# Apolipoprotein E4 has extensive conformational heterogeneity in lipid free and bound forms

**DOI:** 10.1101/2022.02.02.478828

**Authors:** Melissa D. Stuchell-Brereton, Maxwell I. Zimmerman, Justin J. Miller, Upasana L. Mallimadugula, J. Jeremias Incicco, Debjit Roy, Louis G. Smith, Berevan Baban, Gregory T. DeKoster, Carl Frieden, Gregory R. Bowman, Andrea Soranno

**Affiliations:** Department of Biochemistry and Molecular Biophysics, Washington University in St Louis, 660 St Euclid Ave, 63110, Saint Louis, MO, USA; Center for Science and Engineering of Living Cells, Washington University in St Louis, 1 Brookings Drive, 63130, Saint Louis, MO, USA

## Abstract

The ε4-allele variant of Apolipoprotein E (ApoE4) is the strongest genetic risk factor for Alzheimer’s disease, though it only differs from its neutral counterpart ApoE3 by a single amino acid substitution. While ApoE4 influences the formation of plaques and neurofibrillary tangles, the structural determinants of pathogenicity remain undetermined due to limited structural information. We apply a combination of single-molecule spectroscopy and molecular dynamics simulations to construct an atomically-detailed model of monomeric ApoE4 and probe the effect of lipid association. Our data reveal that ApoE4 is far more disordered than previously thought and retains significant conformational heterogeneity after binding lipids. In particular, the behavior of the hinge region and C-terminal domain of ApoE4 differs substantially from that proposed in previous models and provides a crucial foundation for understanding how ApoE4 differs from non-pathogenic and protective variants of the protein.

---

Apolipoprotein E (ApoE) is a 299 amino acid protein involved in lipid-transport and cholesterol homeostasis^1,2^ that plays a key role in Alzheimer’s disease (AD). The polymorphic nature of APOE allows for encoding three variants (ApoE2, ApoE3, ApoE4)^3^ that have dramatic functional differences, even though it is only a single amino acid change that differentiates ApoE3 from ApoE2 (R158C) and ApoE4 (C112R)^4^. The most striking example is ApoE4, which is recognized as the major genetic risk factor for AD^5–9^, with individuals who are homozygous for the ε4-allele having up to fifteen-fold higher probability of developing late onset AD^10,11^. In contrast, ApoE3 appears to have no impact on the progression of AD, while ApoE2 has been proposed to be protective toward the disease^12^. A current hypothesis is that these functional differences stem from structural changes imposed upon ApoE by this single residue substitution, and thus having a potential impact on its interaction with AD factors, such as amyloid-beta plaques, and neurofibrillary tangles^13,14^. In both the cardiovascular and the central nervous system, ApoE is prevalently associated non-covalently with lipids as part of lipoproteins and the single-point mutations are known to alter its interaction with specific lipoprotein populations^15^. From a biochemical point of view, previous work from Garai *et al* suggests that only the monomeric form – not the oligomers – is competent for high-affinity lipid binding^16^. Therefore, understanding the monomeric structure of ApoE is key to unmasking the mechanisms controlling its interaction with lipids. In addition, recent experiments have found that ApoE expressed by microglia and astrocytes can also occur in poorly- and non-lipidated forms^17^. However, a structural characterization of monomeric ApoE in its lipid-free states remains elusive. One major obstacle is posed by the high propensity of ApoE to form oligomers^18^, which hampers the investigation of the monomeric form (see **Supplementary Information**). A second challenge is the disordered nature of numerous short segments of the protein, which have been proposed to be flexible and confer structural heterogeneity^19^ rendering these regions invisible to conventional structural biology methods.

ApoE comprises four different regions: the N-terminal tail (1-23), the four-helix bundle (24-167)^20–22^, the hinge region (168-205), and the C-terminal domain (206-299) (**Fig. 1**). Current conformational models^19,23^ of the monomeric lipid-free ApoE agree on the structure of the four-helix bundle^20–22^, but they disagree on the configurations of the hinge and C-terminal region and their orientation with respect to the N-terminal domain. Ensemble Förster Resonance Energy Transfer (FRET) and EPR studies^24^ suggest ApoE4 forms a close contact between the four-helix bundle and the C-terminal domain, whereas ApoE3 explores more open conformations. This is at odds with the compact set of structures determined by NMR on a monomeric ApoE3-like variant^22^. Recent HDX experiments identified isoform-dependent differences in solvent accessibility of the N-terminal domain, hinting that single-point mutations affect the ability of the C-terminal domain to shield specific regions of the four-helix bundle^19^. However, the interpretation of ensemble FRET, EPR^24^, and HDX experiments^19^ is complicated by the fact that measurements were performed under conditions in which the protein is a stable tetramer^16,19^ and, therefore, are not representative of the conformations of the protein in its monomeric form. The same limitation applies to previous investigations of the folding stability of the protein domains^16,25–27^ and its interaction with lipids^24–29^, where ApoE was investigated at concentrations that favor either dimer or tetramer conformations^16,24,28,29^.

**Figure 1.**
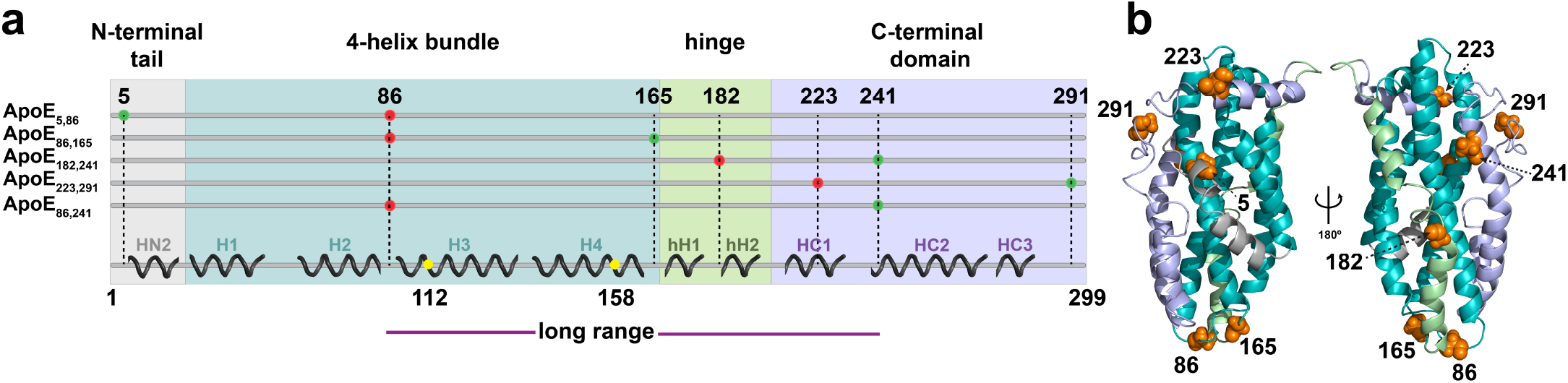
Protein structural regions and single-molecule constructs of full-length ApoE4. **a**. Schematic representation of the secondary structure content in ApoE4 based on the NMR structure (PDB: 2L7B) of the ApoE3-like variant with corresponding designations and identification of the major protein domains: N-terminal tail (gray), four-helix bundle (teal), hinge region (green), and C-terminal domain (light purple). Labeling positions are identified on the linear sequence. Position A86C is located in the random coil between helices H2 and H3 as previously defined ^20,31^ and serves as a common reference point to investigate the folded N-terminal domain from two different perspectives. When paired with position A5C (**ApoE4_5,86_**), which is situated upstream of the start of the H1 helix, A86C monitors the conformational properties and folding stability of the N-terminal tail. When paired with position G165C (**ApoE4_86,165_**), which is located at the end of the H4 helix, A86C provides a read out for the folding of the four-helix bundle^22,31^. Positions G182C and A241C (**ApoE4_182,241_**) allow monitoring the behavior of the hinge domain with respect to the C-terminus, while positions S223C and A291C (**ApoE4_223,291_**) provide information on the structural properties of the C-terminal domain. Finally, probe positions located at A86C and A241C (**ApoE4_86,241_**) allow us to monitor long-range interactions between the N- and C-terminal domains. **b**. 180-degree rotated views of the monomeric ApoE3-like variant NMR Structure (PDB: 2L7B) highlighting labeling positions shown in orange. Structure color differentiates major protein domains described in **a**.

Here, we circumvent these experimental difficulties by harnessing single-molecule fluorescence spectroscopy, an approach that enables working at sufficiently low protein concentrations to avoid oligomerization and directly access the protein in its monomeric form. Single-molecule FRET provides a direct readout on the conformations and stability of specific domains within full-length ApoE4, in both the lipid-free and lipid-bound states. We further complement single-molecule observations with molecular dynamics (MD) simulations to obtain an atomically-detailed representation of protein conformations that is consistent with our experimental data.

## RESULTS

To study the conformations of ApoE4 *via* single-molecule FRET, we designed, expressed, and purified five distinct full-length double-cysteine mutants of the protein (see **Fig. 1a and Supplementary Information**). We used the ApoE3-like structure determined by NMR^22^ (**Fig. 1b**) as a blueprint to guide our choice of labeling positions, such that each dye pair combination probes one of the four regions of the protein.

### Folding and stability of the four-helix bundle

We first focus on the **ApoE4_86,165_** construct, where labeling positions are located in the random coil between helices H2 and H3 (A86C) and at the end of helix H4 (G165C), which enables probing the folding of the four-helix bundle. Although 79 amino acids apart in the sequence, the two labeling positions are expected to be in close proximity with a predicted transfer efficiency of 0.99 (see **Fig. 1b**), based on the ApoE3-like NMR structure^22^. Indeed, under aqueous buffer conditions (50 mM NaPi, pH 7.4), single-molecule FRET measurements of **ApoE4_86,165_** display a narrow distribution of transfer efficiencies with a mean value of 0.98 ± 0.01 (**Fig. 2 and Supplementary Table 1**), compatible with the folded four-helix bundle. With increasing concentrations of Guanidinium Chloride (GdmCl) (**Fig. 2**), the amplitude of the population at high transfer efficiency decreases in favor of two other populations characterized by distinct mean transfer efficiencies. One population is observed at E ~ 0.62 across different GdmCl concentrations and its relative abundance exhibits a non-monotonic trend, increasing between 0 and 1.5 M GdmCl and then decreasing until its disappearance at ~ 3 M GdmCl (**Fig. 3a**), which is consistent with a folding intermediate. The measured lower transfer efficiency, compared to the folded state, is compatible with a more expanded conformation (**Supplementary Fig. 1**), suggesting a partial unpacking of the four-helix bundle. The other population reveals a continuous shift in transfer efficiencies from 0.35 to 0.2 when moving from low to high denaturant concentration **(Fig. 3a)**, which is accompanied by a continuous increase in its relative abundance (**Fig. 3b**). This is consistent with the behavior expected for an unfolded region undergoing denaturation^30^. By fitting the relative abundance of each population with a three-state model, we quantify the stability of the intermediate and folded state, which are ΔG_0_^UI^ = −5.6 ± 0.4 RT and ΔG_0_^UF^ = −8.3 ± 0.4 RT, respectively (**Fig. 3c, Supplementary Fig. 2 and Supplementary Table 2**). The midpoint of the unfolding transition occurs at ~2 M GdmCl (**Fig. 3b**), which is in excellent agreement with previous ensemble experiments^25–27^ (**Supplementary Table 3**).

**Figure 2.**
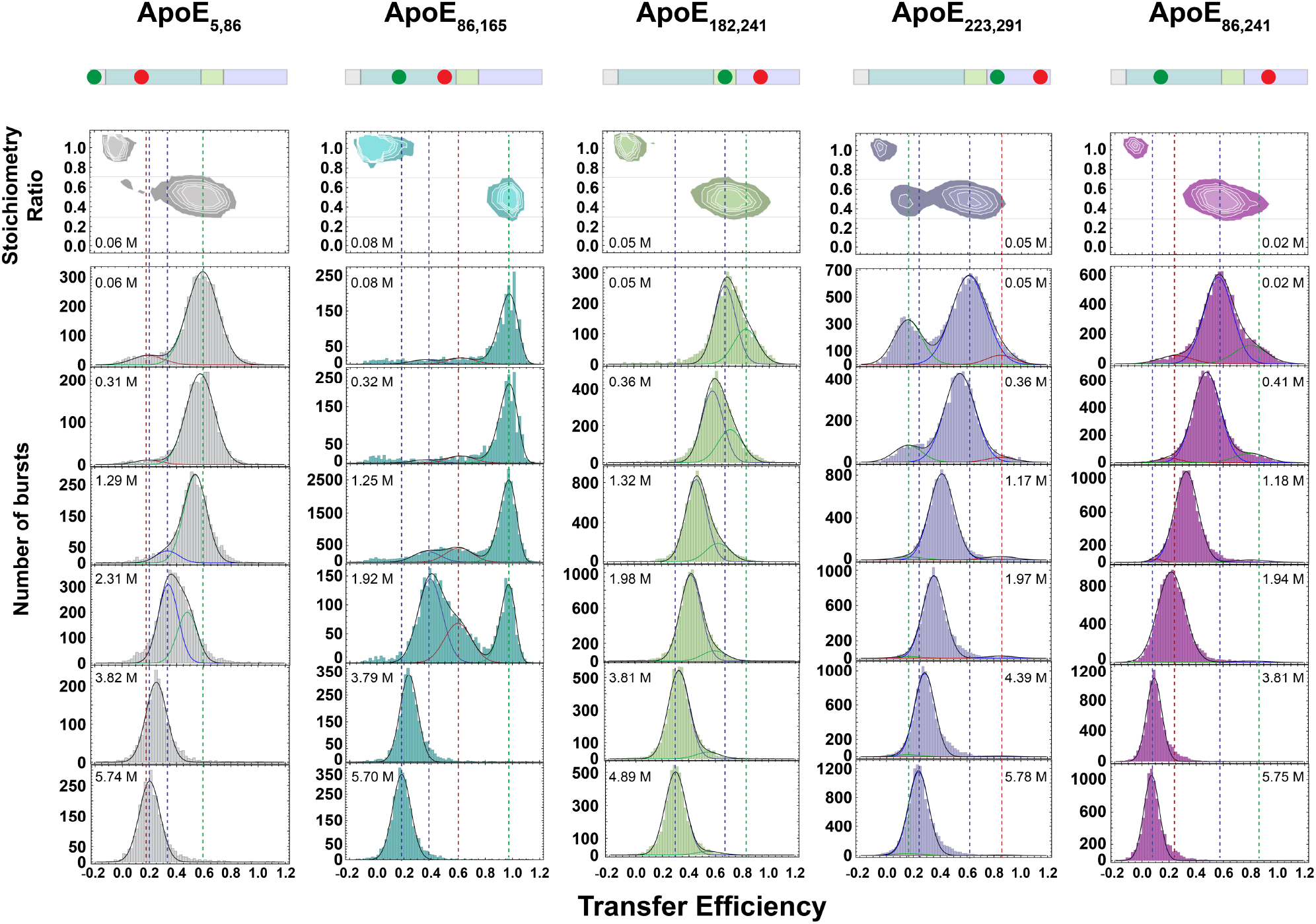
Single-molecule fluorescence experiments of lipid-free full-length ApoE4. Transfer efficiency histograms for selected bursts with fluorescence stoichiometry ratio between 0.3 and 0.7 across the five full-length constructs **ApoE4_5, 86_** (gray), **ApoE4_86, 165_** (teal), **ApoE4_182, 241_** (green), **ApoE_223, 291_** (light purple), **ApoE4_86, 241_** (magenta) at increasing concentrations of GdmCl. Under aqueous conditions all histograms reveal coexistence of multiple states. Lines are visual guides for contrasting the native and completely unfolded configurations in each construct.

**Figure 3.**
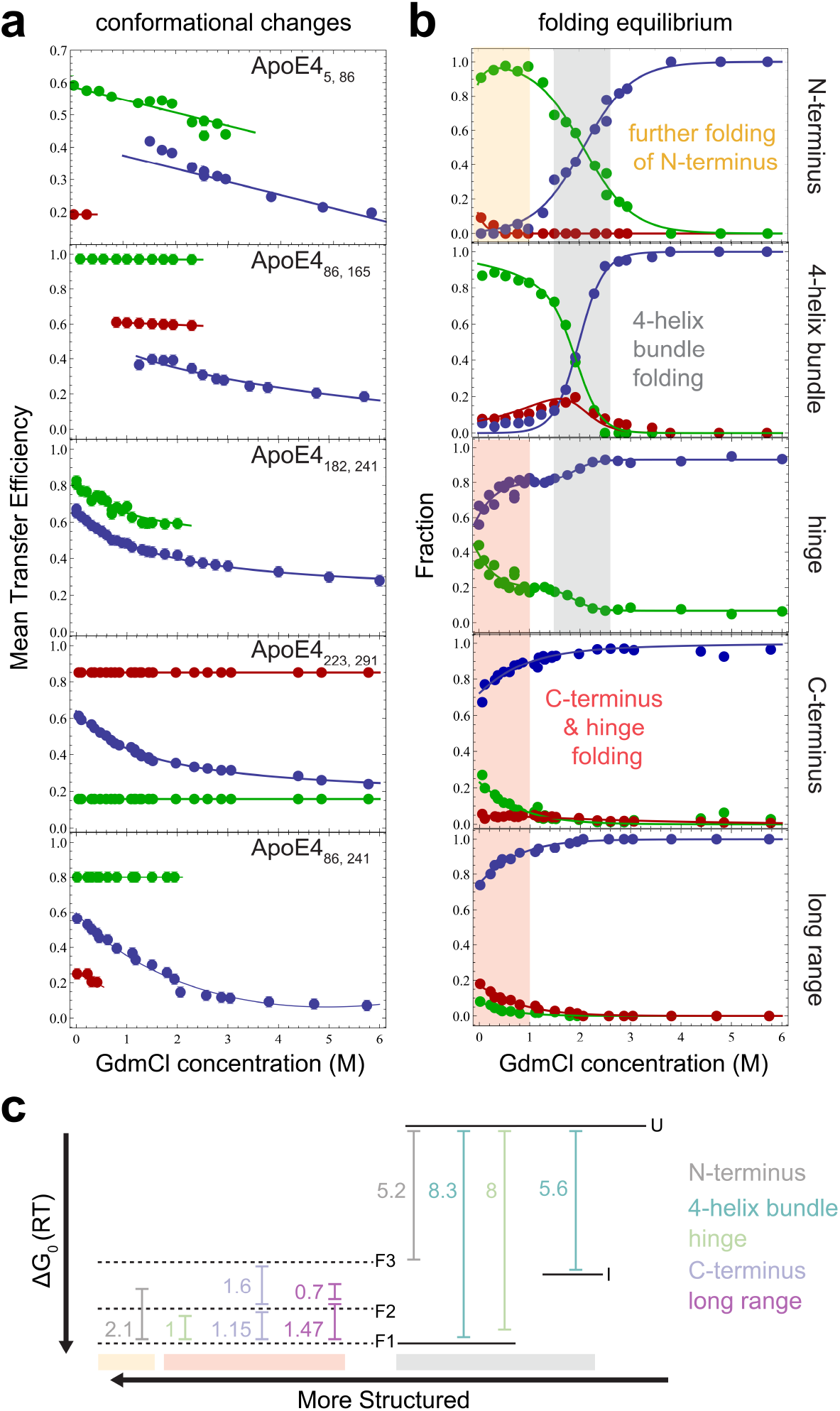
Mean transfer efficiencies and relative fractions of populations for lipid-free ApoE4. **a**. Blue, red, and green identify corresponding populations in transfer efficiency histograms of **Fig. 2**. Solid lines connect mean transfer efficiencies to simply provide a visual guide. Mean transfer efficiencies are shown only for population fractions larger than 10% or when analyzed assuming a fix shared value. Associated standard deviation errors are reported in **Supplementary Table 1**. **b**. Solid lines reflect independent fits with three-state equilibrium between the different conformers (**Supplementary Table 2** and **Supplementary Fig. 2**). Vertical shaded areas indicate folding across specific regions. **c**. Free energy diagram of identified states in the 4-helix bundle, hinge, N- and- C-terminus, and from long range measurements. Solid lines represent the equilibrium between completely unfolded protein (U), formation of the intermediate (I) and complete folding of the 4-helix bundle (F). Dashed lines indicate the different folded states identified in the N-terminal tail, hinge region, C-terminal domain, and long-range contacts. Dashed lines are used to underline that these different configurations coexist with the folded state of the four-helix bundle.

### The N-terminal tail

We complete the investigation of the N-terminal domain by focusing on the N-terminal tail, which is not resolved in the crystal structure of the four-helix bundle^31^. Position A5C is situated upstream of the start of helix H1 and when paired with A86C monitors the conformational properties of the N-terminal tail (**Fig. 1**). Single-molecule FRET measurements of **ApoE4_5,86_** reveal two distinct populations in equilibrium under aqueous buffer conditions. The more abundant population has a mean transfer efficiency of 0.61 ± 0.02, while the less abundant population sits at 0.21 ± 0.05 (**Fig. 2**). Comparing the donor lifetime *vs* transfer efficiency indicates that the population at low transfer efficiency is compatible with a rigid distance where positions 5 and 86 are located ~ 7 nm apart (**Supplementary Fig. 1)**. Conversely, the population at higher transfer efficiency follows the expected trend of a dynamic conformational ensemble, that is, an ensemble of inter-dye distances that are sampled in a time-scale much shorter than the residence time of the protein in the confocal volume. Interestingly, the results are better described using a wormlike chain distribution with persistence length *l_p_* (an estimate of the minimal flexible segment) equal to 2.5 nm and contour length *l_c_* (the maximum extension of the probed region) equal to 7.7 nm (**Supplementary Fig. 1**). Note that this contour length is just ~25% of the contour length expected for an equivalent fully disordered region, suggesting that secondary structure formation occurs within this population. To further test for the presence of secondary structure, we investigated the effect of denaturant. We observe that the population at low transfer efficiency is completely destabilized at 0.5 M GdmCl and that the population at higher transfer efficiency tends to shift towards lower values with increasing denaturant (**Fig. 3a**). This result is consistent with a population that is not completely structured and contains a certain degree of flexibility^30^. Interestingly, an inflexion point in the mean transfer efficiency of this population occurs between 1 and 2 M GdmCl accompanied by a change in the width of the distribution (**Supplementary Fig. 3-4**). We interpret this behavior as the results of the coexistence of two populations with similar transfer efficiencies within the same observed peak. By fitting two independent populations within the mean transfer efficiency distribution (**Fig. 2**), we obtain a midpoint of the transition (c_1/2_) equal to 2.06 ± 0.01 M and a ΔG_0_ equal to 5.2 ± 0.2 RT (**Fig. 3c**, compare alternative analysis **in Supplementary Information**). This observation can be understood considering that positions 5 and 86 sample not only the N-terminal tail but also helix H1 and H2 of the four-helix bundle.

### The hinge region

Positions G182C and A241C (**ApoE4_182,241_**) allow monitoring of the behavior of the hinge domain with respect to the C-terminus. Analysis of the corresponding transfer efficiency histograms reveals an asymmetric distribution of transfer efficiencies under aqueous buffer conditions. We analyze the asymmetric distribution in terms of two distinct populations (**Fig. 2**). The population associated with lower mean transfer efficiency (E = 0.62 ± 0.02) accounts for 60% of the observed molecules, whereas the high transfer efficiency population (E = 0.83 ± 0.02) accounts for the remaining 40%, corresponding to a free energy difference between these states of 1.0 ± 0.2 RT (**Supplementary Table 1-2**).The asymmetry of the distribution persists with increasing denaturant concentrations, with both populations shifting toward lower transfer efficiencies **(Fig. 3)**, as expected for disordered or partially disordered regions^30^. Comparing lifetime and transfer efficiency indicates that both populations reflect dynamic averages that, similarly to the case of the N-terminal tail, we can describe in terms of a wormlike chain (**Supplementary Fig. 1**). Interestingly, the relative abundance of the two populations reveals a second transition in the range between 1.5 and 2.5 M GdmCl concentration. The range of this transition coincides with the same range observed for the folding transition of the four-helix bundle (c_1/2_ = 1.9 ± 0.2 M, **Supplementary Table 2**) and suggests a conformational change of the hinge region contextually with the folding of the N-terminal domain.

### The C-terminal domain

Positions S223C and A291C (**ApoE4_223,291_**) provide information on the structural properties of the C-terminal domain. Under aqueous buffer conditions, we observe a broad distribution of transfer efficiencies that correspond to at least three distinct conformational states sampling long-, middle-, and shortrange distances between the fluorophores **(Fig. 2)**. When comparing donor lifetime and transfer efficiency, the population at 0.13 ± 0.04 mean transfer efficiency is compatible with a rigid region of ~7.9 nm (**Supplementary Fig. 1)**. This population accounts for 27 ± 4% of the protein configurations and is completely destabilized in favor of the other populations above 1.2 M GdmCl. Different is the case for the population with transfer efficiency equal to 0.61 ± 0.02, whose donor lifetime follows the expected trend for a dynamic ensemble and whose relative abundance is stabilized by increasing concentrations of denaturant. Both elements point toward a population that is more flexible and, at least, partially disordered, as further supported by the continuous shift of the peak from high to low transfer efficiencies when tuning the solvent quality from a poorer solvent (aqueous buffer) to a better solvent (GdmCl). The increased broadening of the width of this population below 1 M GdmCl (**Supplementary Fig. 3**), which exceeds the width measured for other constructs, points to an increased heterogeneity due to structure formation. This is consistent with previous characterizations of the C-terminal region, where destabilization of the secondary structure was observed above 1 M GdmCl^25,26^. The third population at ~ 0.85 mean transfer efficiency represents more compact configurations of the C-terminal domain, where position 223 and 291 are brought in close proximity. Interestingly, the small relative abundance of this population decreases above 1 M GdmCl and disappears at 2.75 M GdmCl (**Fig. 3**). This regime of concentrations coincides with the folding of the four-helix bundle and mirrors that observed for the hinge region, suggesting that folding of the four-helix bundle induces conformational changes in the C-terminal region.

### Long-range interactions

To better understand whether the interactions between the four-helix bundle and the C-terminal region are due to stable contacts or reflect long range electrostatic interactions, we further investigate the transfer efficiency distribution between A86C and A241C (**ApoE4_86,241_**). Under aqueous buffer conditions, we observe the occurrence of at least three populations with corresponding mean transfer efficiencies of 0.24 ± 0.01, 0.59 ± 0.02, and 0.87 ± 0.02 (**Fig. 2**). This is consistent with observation of multiple configurations in both the hinge and C-terminal regions. When comparing donor lifetime and transfer efficiency (**Supplementary Fig. 1**), all the populations lie on the trend expected for a dynamic ensemble, suggesting that the observed conformations in the hinge and C-terminus arise by means of long-range forces with the four-helix bundle. The expanded population is destabilized at very low GdmCl concentrations, suggesting the interaction between the domains can be easily disrupted by competing interactions, such as ion screening of the electrostatic forces. Interestingly, a small percentage of the collapsed state persists up to concentrations of denaturant that are compatible with the unfolding of the N-terminal domain.

### MD simulations confirm structural heterogeneity

To gain insights into the structural details of the conformational ensemble of ApoE4, we performed all-atom MD simulations of the full-length protein on the distributed computing platform Folding@home for a total aggregated time of 3.45 ms. We then constructed a Markov State Model to bin the conformational ensemble into unique states. For each observed state, we modeled fluorophores onto the labeling positions *post-hoc* and reconstructed a set of transfer efficiency histograms that accounts for shot noise and the kinetic averaging of conformations in the observation timescale **(see Supplementary Information)**. The comparison between simulated and measured transfer efficiency histograms is shown in **Fig. 4a**. We find good agreement between both data sets, including the occurrence of a multimodal transfer efficiency distribution for **ApoE_223,291_**. To better disentangle the conformations underlying the simulated transfer efficiency histograms, we analyzed the simulation data for the occurrence of correlations across all distance pairs (**Fig. 4b and Supplementary Fig. 5**). This analysis reveals three subpopulations associated with the distance between positions 86 and 165 whose mean transfer efficiencies fall within the observed distribution for **ApoE_86,165_**. The conformational changes in these subpopulations are not restricted to these specific labeling positions but propagate across the entire protein, highlighting correlated changes in the hinge region and anticorrelated ones in the C-terminal domain. In particular, the identified subpopulations in each distance pair correlation parallel the distance and relative abundance trends observed in the experiments. All three identified subpopulations differ from the ApoE3-like NMR structure, where numerous contacts previously identified between the four-helix bundle and the C-terminal domain are not observed even in the more compact conformations. (**Fig. 5**). Alignment of subpopulation structures reveals how these correlative trends reflect different degrees of conformational heterogeneity in the protein (**Fig. 4c**). We refer to the three major subpopulations as *closed*, where the C-terminal domain is docked on the four-helix bundle, *open*, where the C-terminal domain is undocked, and *extended*, where the undocked C-terminal domain adopts more extended configurations. Interestingly, these conformational differences do not stem from varying degrees of secondary structure in the C-terminal domain (see **Supplementary Fig. 6**). We further analyzed the simulations to verify whether specific residue contacts are maintained despite the extensive conformational heterogeneity. We identified a set of persistent contacts within the four-helix bundle and two additional contacts between the four-helix bundle and the HC1 helix of the C-terminal domain, which suggests that the relative position of HC1 with respect to the four-helix bundle is maintained across all the subpopulations. (**Fig. 5–6**). At variance with the *closed* subpopulation, the *open* and *extended* ensembles show an increase in the number of contacts of the N-terminal tail with the four-helix bundle and the HC1 helix, which may dictate whether the C-terminal domain docks onto the four-helix bundle. Interestingly, there are no shared contacts across the three subpopulations within the N-terminal tail or the hinge region (**Fig. 6**, highlighted in yellow), which suggests that these regions are adopting different conformations in each state. Indeed, the position of the hinge region differs across the three subpopulations and is directed by interactions between the hH1 helix and either the N-terminal tail or the four-helix bundle (**Fig. 6**). Specifically, in the *closed* configuration the hinge region mainly interacts with helices H1 and H2, whereas in the *open* and *extended* configurations the hinge explores the surface of helices H2 and H3 with differing extent of specificity. Altogether, MD simulations confirm the experimental observation that lipid-free ApoE4 adopts a dynamic structural ensemble with at least three distinct states.

**Figure 4.**
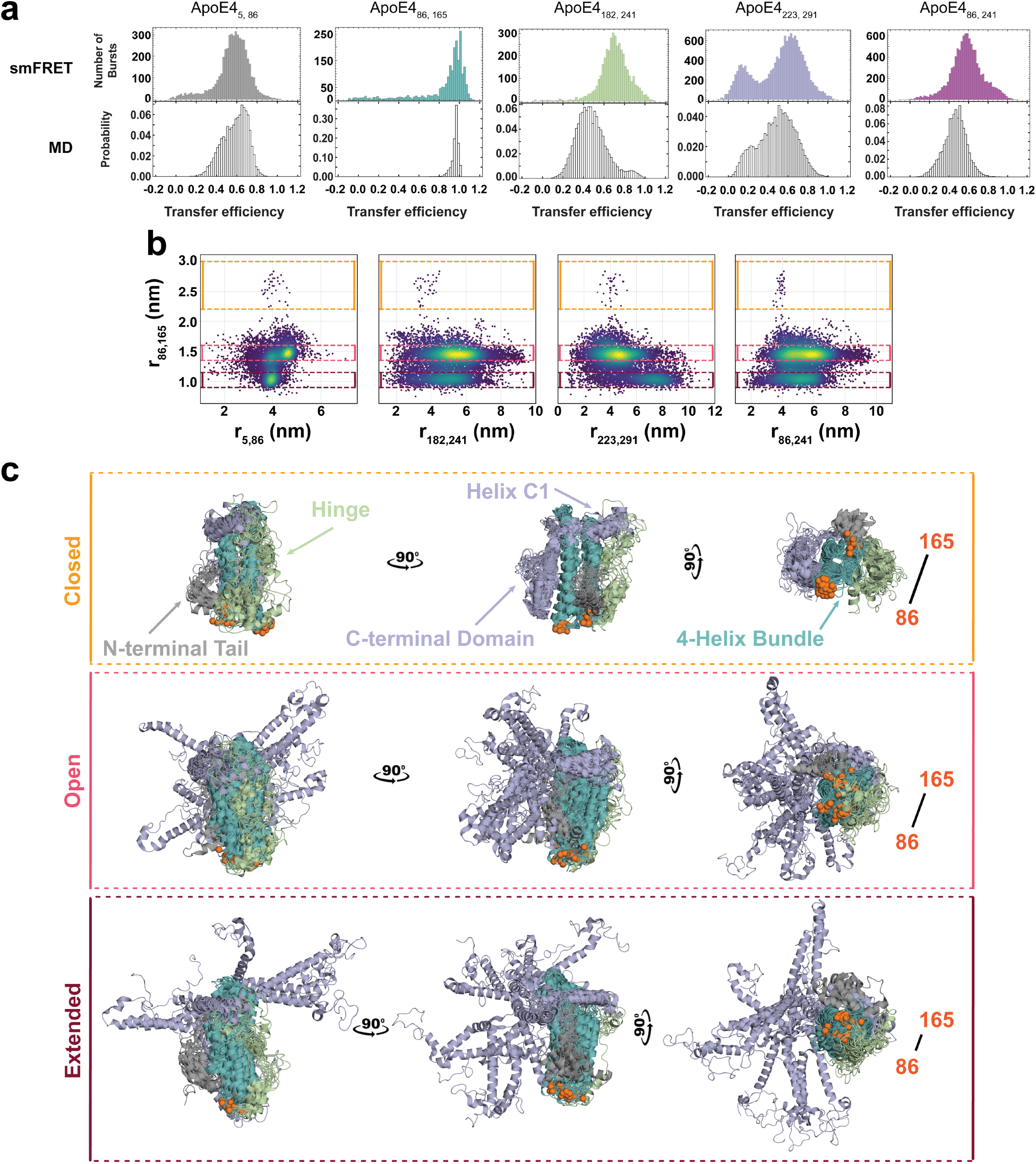
Comparison between transfer efficiency histograms in single-molecule measurements and MD simulations for lipid-free ApoE4. **a**. Single-molecule FRET histograms of the five investigated constructs **ApoE4_5, 86_** (gray), **ApoE4_86, 165_** (teal), **ApoE4_182, 241_** (green), **ApoE4_223, 291_** (light purple), **ApoE4_86, 241_** (purple) are compared with equivalent distribution of transfer efficiencies computed from MD simulations (white). **b**. Distance pair correlations from MD simulations contrasting the distance **r_86, 165_** with the distances **r_5,86_, r_182,241_**, **r_223,291_, r_86,241_**. Colored boxes (yellow, red, brown) identify three major configuration regimes of the four-helix bundle and corresponding changes in the other protein regions. **c**. The fifteen most probable configurations for each of the three states closed, open, and extended, as identified from the data in panel **b**. Position of 86 and 165 fluorophores is highlighted in orange, whereas the N-terminal tail is displayed in gray, the four-helix bundle in teal, the hinge region in green, and C-terminal domain in light purple (compare with **Fig. 1**).

**Figure 5.**
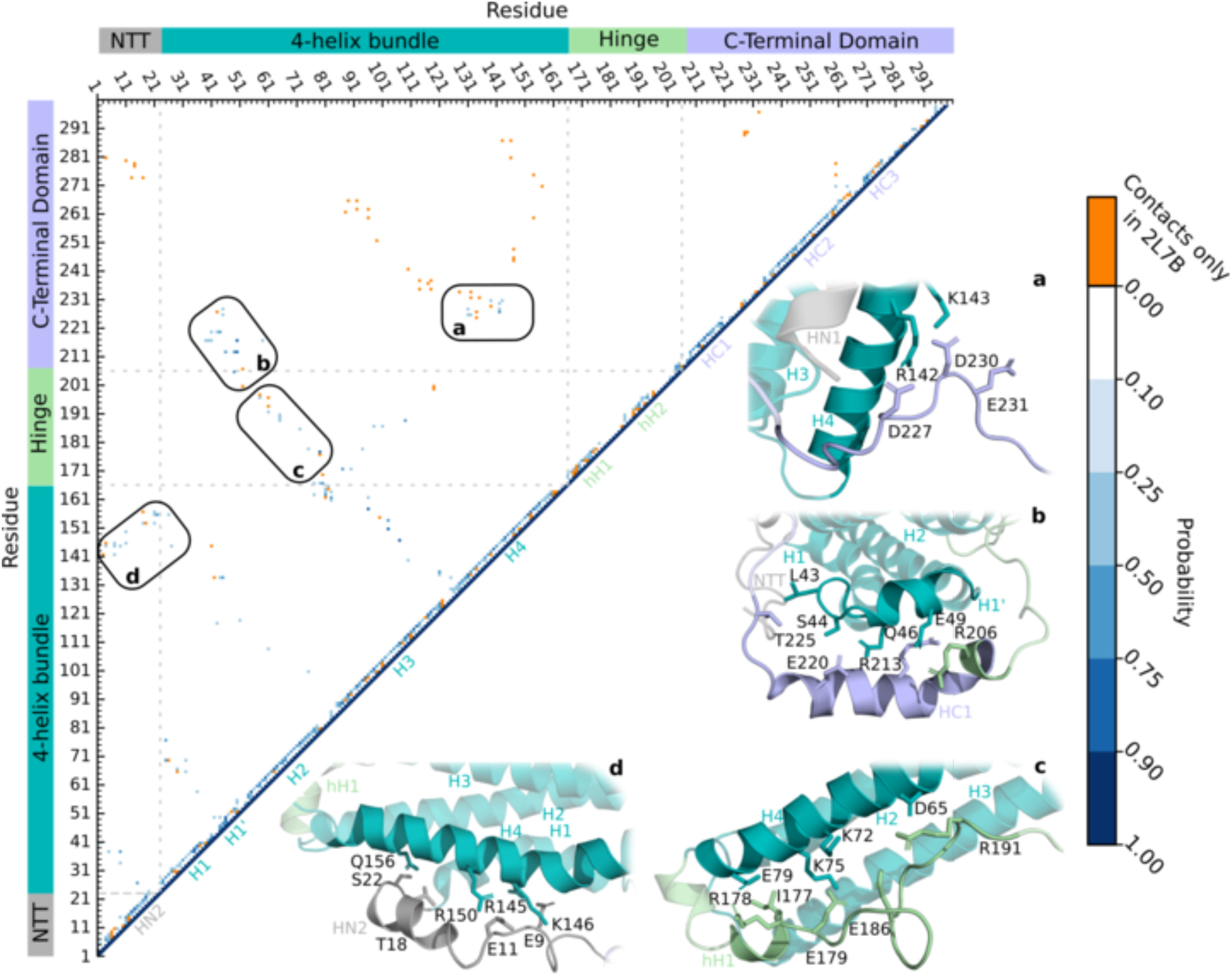
Contacts in lipid-free ApoE4. Probability contact map from all structures from MD simulations contrasted against contacts found in the ApoE3-like NMR structure (PDB:2L7B). Contacts that are present only in the NMR structure are reported in orange. Black boxes identify contacts that are due to domain-domain interactions.

**Figure 6.**
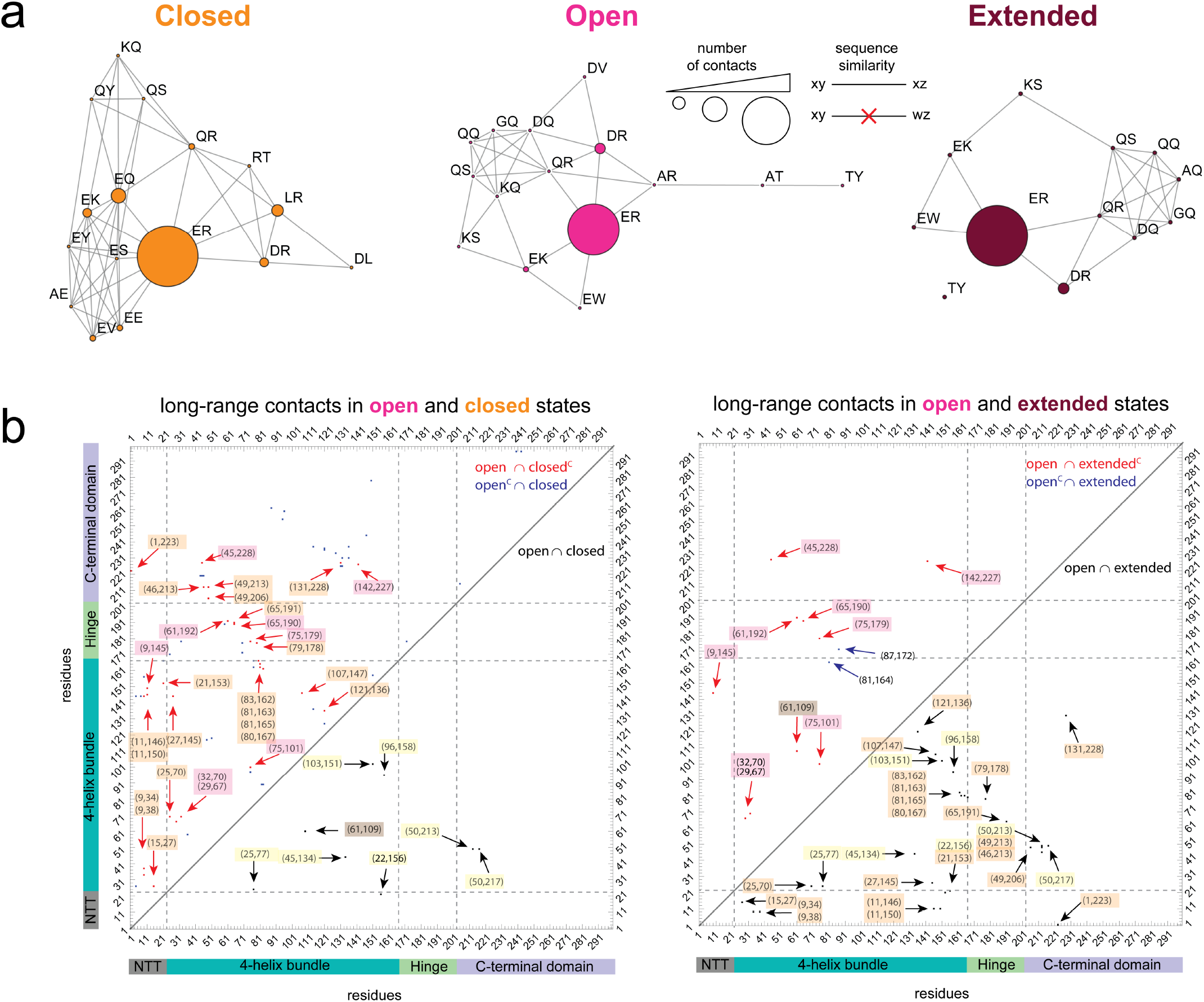
Long-range contact differences across the *closed, open*, and *extended* subpopulations of ApoE4. Long-range contacts here are identified residues whose centers of mass are less than 3 Å apart from each other and that are separated in sequence by at least 6 residues. **a**. Interacting residues identified in the *closed, open*, and *extended* subpopulations. Letters represent amino acid codes. Nodes are scaled according to the number of contacts and edges connect contacts based on sequence similarity. The majority of contacts occurs between charged residues (e.g., glutamic acid and arginine). **b**. List of long-range contacts. *Left panel*: contacts that are in the *open* and *closed* configuration (open ∩ closed, black), contacts that are in the *open* but not in the *closed* configuration (open ∩ closedC, red), and contacts that are in the *closed* but not *open* configuration (open^C^ ∩ closed, blue). *Right panel*: contacts that are in the *open* and *extended* configuration (open ∩ extended, black), contacts that are in the *open* but not in the *extended* (open ∩ extendedC, red), and contacts that are in the *extended* but not in the *open* (openC ∩ extended, blue). Highlighted in yellow: contacts that are shared across all three states (closed ∩ open ∩ extended). Highlighted in orange: contacts that are in the *open* and *extended*, but not in the *closed* configurations (open ∩ extended ∩ closedC). Highlighted in red: contacts that are in the *open* but not in the *extended* and not in the *closed* configurations (open ∩ extended^C^ ∩ closed^C^). Highlighted in brown: contacts that are in both *open* and *closed*, but not in the *extended* configuration (open ∩ closed ∩ extendedC).

### Lipid association of ApoE4

Finally, we turn to investigating how the structural heterogeneity of ApoE4 is impacted by binding to lipids, which reflects the most likely populated configuration under physiological conditions. To this end, we focus on the interaction between the ApoE4 constructs and Dipalmitoylphosphatidylcholine (DMPC) liposomes with an average radius of 40 ± 20 nm (**Fig. 7a** and **Supplementary Fig. 7**). We chose DMPC because it is a good mimic of the lipids found in lipoproteins, both in terms of hydrophilic head group and average length of the fatty acid chain^32,33^. Using single-molecule FRET and a high concentration of liposomes (100 μg/ml), we tested whether the labeled constructs could bind to lipids. **ApoE4_5,86_** (N-terminal tail), **ApoE4_223,291_** (C-terminal domain), and **ApoE4_86,241_** (long-range contacts) all exhibit a single narrow distribution of transfer efficiencies with a clear shift of the mean toward values lower than 0.2, representing very extended states of the protein (**Fig. 7b,e,f** and **Supplementary Fig. 8**). The complete disappearance of the populations observed for lipid-free ApoE4 confirms that these three constructs are fully associated with lipids. Interestingly, the construct **ApoE4_86,165_** (four-helix bundle) exhibits two coexisting populations in equilibrium, one at high transfer efficiency (0.894 ± 0.004) and one at low transfer efficiency (0.037 ± 0.006) (**Fig. 7c** and **Supplementary Table 4**). Neither transfer efficiency is compatible with the population measured in aqueous conditions in the absence of lipids. This suggests that the four-helix bundle can undergo unpacking and restructuring when associated with lipids and that a certain degree of heterogeneity, represented by these two distributions of transfer efficiencies, is conserved even in the lipid-bound state (**Fig. 7g-h**). Finally, the **ApoE4_182,241_** (hinge region) construct also supports the occurrence of at least two distinct configurations of ApoE4 in the lipid-bound state (**Fig. 7d**), though the relative ratio between the two bound states is different compared to **ApoE4_86,165_**. This observation further reflects how the hinge and N-terminal domains are interconnected regions that maintain a certain degree of independence. Overall, taken together, these data support that the protein is completely associated with lipids at the studied concentration. We further analyzed the change in the fluorescence stoichiometry ratio of the lipid-bound vs lipid-free conformations for each construct and validate that we are observing one single protein per lipid-bound state. Binding of multiple proteins in the lipid-bound state would result in a significant change in stoichiometry since a non-negligible fraction of molecules is double-labeled with only acceptor or donor fluorophores. The negligible variation in fluorescence stoichiometry suggests that the protein is monomeric (**Supplementary Fig. 9**). Finally, we observed that the low transfer efficiencies measured for lipid-bound ApoE correspond to relatively short distances (< 10 nm) (see **Supplementary Fig. 8**) when compared with the liposome size, posing the question on whether the protein is bound to the liposome or some portion of the liposome. Correlating the fluorescence signal (either from donor or acceptor direct excitation) in the same single-molecule measurements, we quantified the size of the lipid-bound states. The measurements clearly reveal an increase in the hydrodynamic radius of approximately 2-3 times the dimension of the lipid-free protein (**Supplementary Fig. 10-11**), which has no overlap with the liposome distribution. Overall, this suggests that during the interaction with liposomes the protein not only undergoes a partial refolding of its domains but does also extract lipids from the larger liposomes in order to create smaller lipid-protein particles (**Fig. 7**).

**Figure 7.**
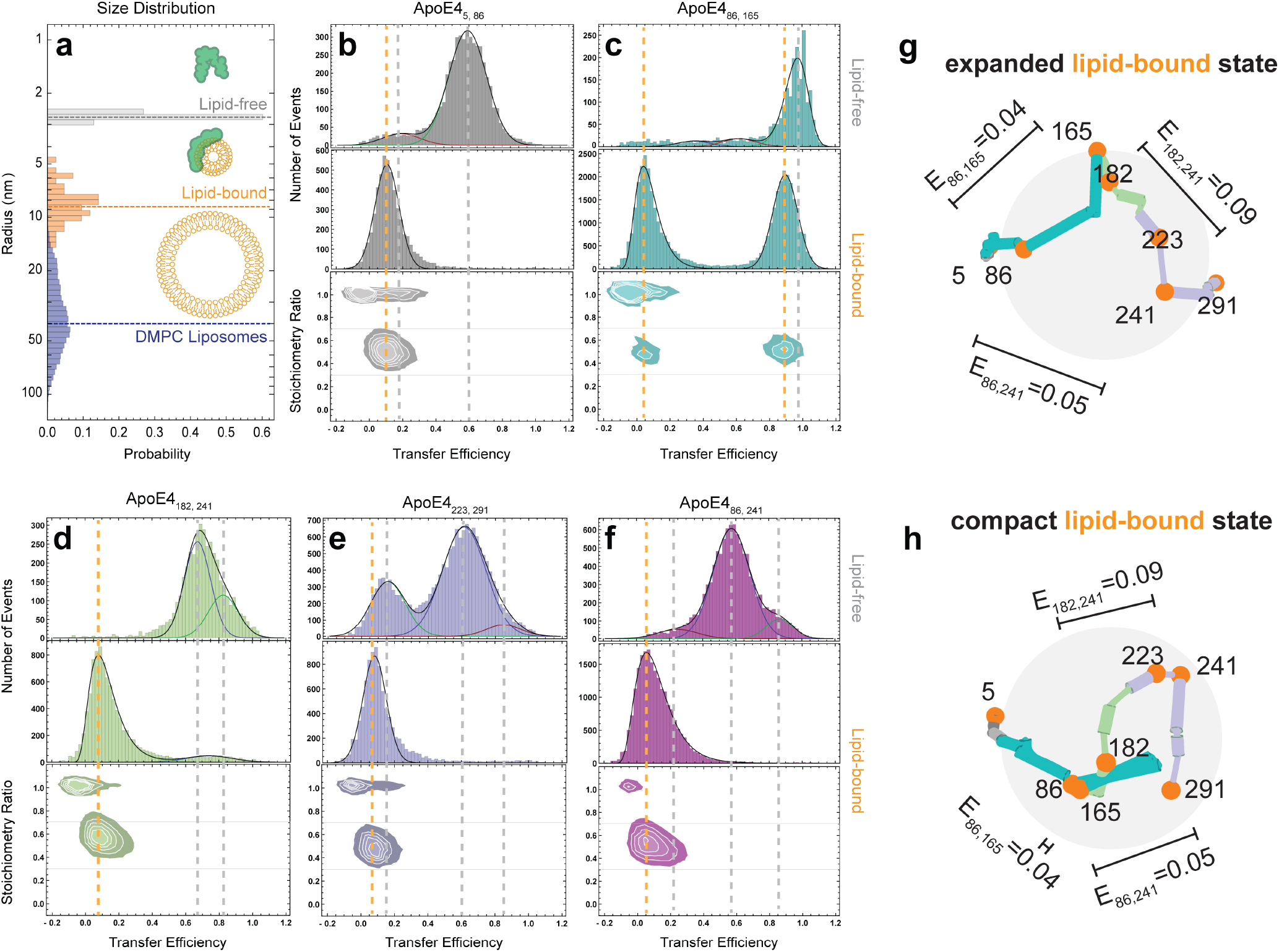
Single-molecule fluorescence experiments on lipid-bound ApoE4. **a**. Distribution of radii for lipid-free ApoE4 (gray, df-FCS), lipid-bound ApoE4 (orange, FCS), and extruded DMPC liposomes (blue, cryo-TEM). **b-f**. Comparison of transfer efficiency histograms for lipid-free and lipid-bound ApoE4 constructs. All histograms report on fluorescent species with a labeling stoichiometry ratio of 1D:1A. Lines are visual guides between the lipid-free (gray) and lipid-bound (orange) mean transfer efficiencies. **g-h**. Representatives examples of conformations of ApoE4 in the expanded and compact lipid-bound states based on an ultra-coarse grained model that satisfies the mean transfer efficiency constraints.

## DISCUSSION

### Conformational heterogeneity in lipid-free ApoE4

Our single-molecule experiments and MD simulations clearly reveal that ApoE4 does not adopt a single structure but, instead, explores a complex and dynamic conformational ensemble. Using the ApoE3-like structure as a reference^22^, we observe large deviations in the conformations of the hinge and C-terminal domain of the protein as well as dynamic fluctuations in the four-helix bundle (**Fig. 4c**). Previous experiments proposed a close proximity of residues 76 and 241^24^ as well as a salt bridge between residues 61 and 255^15,31^, leading to a model where the N- and C-terminal domains are oriented differently to that observed in the NMR structure. Interestingly, in our study we do not observe close contacts of these residues (**Fig. 5–6 and Supplementary Fig. 12-13**) and the identified configurations support the orientation of the N- and C-terminal domains as observed in the ApoE3-like structure. This can be rationalized by noting that experiments that identified these close contacts^24^ were performed under conditions where the protein exists as a dimer or tetramer and therefore may be specific only to these forms of the protein. Similarly, salt bridges^15,31^ have been tested via mutational analysis in the context of lipoproteins or non-monomeric forms of the protein and may reflect other interactions at play. The divergence from the NMR structure (**Fig. 5 and Supplementary Fig. 12-13**) further supports that the key point-mutation C112R that distinguishes ApoE3 from ApoE4 may alter the delicate balance between specific conformers in the structural ensemble, favoring a *closed* configuration in ApoE3. Indeed, our data suggests that the hinge region competes with the C-terminal domain for interactions with the four-helix bundle, where specific contacts involving the N-terminal tail and the four-helix bundle can sway the preference of interaction for one region or the other (**Fig. 6**).

### Folding equilibrium of lipid free ApoE4

Our single-molecule experiments also enable a direct quantification of the stability associated with each conformer of the monomeric protein and provide insights on the overall folding reaction. The denaturant titration suggests that structuring of the N-terminal domain proceeds from a completely unfolded state through an intermediate state where helices H1-H4 are partially formed, followed by the subsequent packing and stabilization of the bundle (**Fig. 3c**). Observation of an intermediate configuration in the four-helix bundle confirms previous interpretation of ensemble data where an intermediate state was presumed^26^. Contextually to the folding of the four-helix bundle, a perturbation occurs in the configurations of the hinge and in the N- and C-terminal tails. While folding of these domains remains largely independent, our data suggests that their structural organization is not disconnected. Indeed, even for labeling positions that do not sample the four-helix bundle, we identify transitions with a mid-point at approximately 2 M GdmCl accompanied by a similar change in free energy (from 5 to 7 RT, see **Fig. 3c** and **Supplementary Fig. 2**). While folding of the C-terminal region is only captured by broadening of the distribution of transfer efficiencies (**Supplementary Fig. 3**), the observation of distinct populations in the N- and C-terminal tails and hinge region provide quantification of the energy difference between these distinct states. The similarity in the relative populations between the hinge and C-terminal regions (as measured by **ApoE4_182,241_** and **ApoE4_223,291_**) across different denaturant concentrations and the overlap between the sequence of the two regions suggests we are monitoring the same configurational change. Therefore, the emerging picture is of a folded four-helix bundle in equilibrium with at least three distinct populations of the C-terminal domain.

### Monomeric ApoE4 forms heterogeneous complexes with lipids

Early EPR studies of ApoE4 suggested that helices in the N- and C-terminal domains remain in close contact in the lipid-bound state, whereas the four-helix bundle undergoes structural rearrangements^28^. A competing model proposed that lipid binding favors a separation between the N- and C-terminal halves of the protein, based on the ApoE3-like NMR structure^19,22^. Interestingly, our data indicates that such an open configuration is a constitutive state explored by the ApoE4 monomer and, therefore, does not require interaction with the lipids to occur. The *open* and *extended* configurations expose the required surface of the C-terminal domain making interaction with lipids possible (**Supplementary Fig. 6a**). Indeed, the region between positions 165 and 270 has been identified as containing Class A amphipathic helices, which can promote lipid binding^34^. Therefore, modulation of the abundance of the open state may impact the affinity of ApoE variants for lipids. Our measurements further indicate that monomeric ApoE can extract lipids and form smaller particles compared to the initial liposome preparation. This observation is compatible with previous measurements monitoring decrease of turbidity in liposome solutions upon addition of ApoE^35–37^. The ability to extract lipids implies an intercalation of the amphipathic helices of the protein within the lipid bilayer. Indeed, amphipathic helices are known to play a key role in non-enzymatic membrane fission^38^, where the membrane fission can be self-propelled by insertion of a first helix that favors insertion of subsequent helices^39–41^. This same mechanism may be at play in the interaction of ApoE with liposomes, where insertion of the C-terminus can then propagate through the hinge to the N-terminus^22^. This model explains how the hinge region, which locks the N-terminal domain in the four-helix bundle structure, can be displaced, leading to a rearrangement of the helices of the bundle and allowing for more expanded configurations. Our experiments indicate that the N-terminal domain adopts at least two different configurations, one where the helices H3 and H4 are in close proximity to one another and one in which the four helices are spread apart on the lipid particle (**Fig.7g-h**). This interpretation is fully compatible with the configurations identified by Henry *et al* ^29^ using crosslinking, mass spectrometry, and simulations of ApoE4, though our data suggests a more expanded configuration of the N-terminal tail (as measured by **ApoE4_5,86_**) and a larger separation between the N- and C-terminal halves of the protein (as measured by **ApoE4_86,241_**). Interestingly, previous simulations of ApoE3 identifies only a close configuration for helices H3 and H4, possibly suggesting a different structural organization of the two variants in their monomeric lipid-bound form^42^.

### Conclusions

The realization that ApoE4 does not adopt one single stable structure, but an intricate conformational ensemble opens the door to new explanations for the mechanism of function of the protein and its role in the context of AD. Our results demonstrate the potential of single-molecule approaches for investigating the relationship between structural ensemble and function of monomeric ApoE. This approach bypasses experimental complications due to protein oligomerization, setting the stage for exploring the impact of sequence variations and interaction with AD factors. Understanding how and why sequence mutations and environmental factors tune ApoE from being a risk factor to having neutral effects is key to identifying appropriate therapeutic strategies that can slow down or even arrest the progression of AD.

## Online Methods

### Protein expression, purification, and labeling

All ApoE4 constructs were expressed in BL21-Gold (DE3) cells (Agilent). The Thioredoxin-His6-ApoE protein fusion was purified using a HisTrap FF column (Cytiva). The tag was cleaved by HRV 3C protease and separated from ApoE4 using a Heparin Sepharose FF column (Cytiva). Anion exchange chromatography (Q Sepharose HP FF column, Cytiva) was then used as the final polishing step. Correct mass of the constructs was analyzed using SDS-PAGE and/or electrospray ionization mass spectrometry (LC-MS). All constructs have been labeled with Alexa 488 and Alexa 594, which serve as donor and acceptor, respectively. For further details see **Supplementary Information**.

### Single-molecule measurements

All single-molecule fluorescence measurements were performed on a Picoquant MT200 instrument (Picoquant, Germany). Single-molecule FRET and Fluorescence Correlation Spectroscopy (FCS) were performed with labeled protein concentrations of 100 pM, estimated from dilutions of samples with known concentration based on absorbance measurements. All single-molecule measurements were performed in 50 mM NaPi pH 7.4, 200 mM β-mercaptoethanol (for photoprotection), 0.001% Tween 20 (for surface passivation) and GdmCl or TMAO (Trimethylamine N oxide) at the reported concentrations, at a room temperature of 295 ± 0.5 K. Pulsed interleaved excitation (PIE) was used to ensure that each burst represents the transfer efficiency determined from a 1:1 donor:acceptor stoichiometry. Importantly, attachment of the probes across different labeling positions has a small impact on the overall protein conformations as measured by dual-focus FCS, which reveals variations across the different constructs of less than 10%. All data were analyzed using the Mathematica package “Fretica” (https://schuler.bioc.uzh.ch/wp-content/uploads/2020/09/Fretica20200915.zip) developed by Daniel Nettels and Ben Schuler. Fluorescence lifetimes (**Supplementary Fig. 14**) are analyzed using a convolution with the Instrument Response Function (IRF) (**Supplementary Fig. 15**). Comparing transfer efficiency estimates from donor lifetimes (reporting about the nanosecond timescale) and from bursts of photons (reporting on the millisecond timescale) enables distinguishing whether the associated population represents a rigid configuration or a dynamic ensemble. In the case of a rigid configuration, the same transfer efficiency is recovered on both timescales and results in a constant value that follows the linear dependence of the lifetime on the mean transfer efficiency. In the case of a dynamic ensemble, a deviation from the linear dependence occurs, which depends on the sampled conformational distribution^30^. For further details see **Supplementary Information**.

### MD simulations

The NMR structure of ApoE3 (PDB ID: 2L7B) was used as a starting point for our simulations, with mutations performed in PyMOL to achieve the structures of ApoE4. We performed 20 rounds of directed sampling harnessing the FAST algorithm^43^ to explore the conformational space of ApoE4, using the residue pairs: R92 and S263, G182 and A241, and S223 and A291, as a directed metric. The resulting simulations were clustered to a shared state space with RMSD of 3.5Å into a total of 18,182 structures that represented the diversity of states explored in our simulations. Each structure was solvated in a dodecahedron box with edges 1.0 nm longer than the largest structure observed in our FAST simulations. Subsequent simulations were launched from these states on the distributed computing platform, Folding@home with 5 independent simulations starting from each state. Each trajectory ran for a maximum of 100 ns, in total reaching an aggregate time of 3.45 ms. Simulations were clustered using distance-based clustering for 15 residue pairs distributed throughout ApoE (5 FRET pairs plus 10 additional residue pairs). Markov State Model was subsequently generated using a lag time of 10 ns and Enspara’s MSMBuilder. Simulations were performed using the Amber03 force-field in combination with TIP3P water model. FAST simulations were performed using GROMACS and Folding@home simulations were performed using OpenMM. FRET histograms were calculated using the smFRET tool deployed in ENSPARA using a rescaling time factor of 225 (see **Supplementary Fig. 16**). For further details see **Supplementary Information**.

## Data, materials, and code availability

MD simulation results will be made available after publication. Main experimental data are available in the Supplementary Tables and raw data will be provided upon request. Plasmid of created constructs will be provided upon request. Code for analysis of single-molecule and computational data is publicly available through the sources indicated in the corresponding sections in the **Online Methods** or **Supplementary Information**.

## Acknowledgments

We thank Alessandro Borgia, Gojun Bu, Anil Cashikar, Hagen Hofmann, Alex Holehouse, David Holtzman, Rohit Pappu, Janice Robertson, and Ben Schuler for helpful discussions. We thank Jasmine Cubuk for assistance with data collection and sample purification. We thank the citizen scientists that donated their computing power through Folding@home to make this work possible. We also thank members of the community that have volunteered their time for technical support and recruitment to the Folding@home community.

This work was supported by the National Institute on Aging (NIH) through P50AG05681-35 (A.S.), R01AG062837 (A.S. and C.F.), U19AG069701 (Project 1: A.S. and C.F.; Core B: G.R.B.) and RF1AG067194 (G.R.B. and A.S.). The content is solely responsibility of the authors and does not necessarily represent the official views of the National Institutes of Health.

A.S. was also funded by the Ruth K Broad Biomedical Research Foundation Extramural Award, the Alzheimer’s Association AARG-591858, and the American Federation for Aging Research RAG18224. C.F. was also funded by the Bright Focus Foundation (A2020382S Award). G.R.B. was also funded by a Packard Fellowship for Science and Engineering from The David & Lucile Packard Foundation and NSF CAREER Award MCB-1552471. J.J.M. was also funded by the NIH training grant T32AG05851804.

## Contributions

M.D.S.B. and A.S. conceived the project and designed the experiments. M.D.S.B. expressed, purified, labeled all the constructs, and performed all single-molecule FRET and FCS measurements. M.D.S.B. and A.S. analyzed all single-molecule measurements. M.Z. performed FAST simulations and developed the FRET histogram algorithm. J.J.M. performed MD simulations on Folding@home. J.J.M., U.M., L.G.S., and M.Z. analyzed the MD simulation data and M.D.S.B. and A.S. contributed to the interpretation. A.S. performed and analyzed ultra-coarse grained MonteCarlo simulations of lipid-bound ApoE. D.R. performed and analyzed TEM experiments. J.J.I. performed and analyzed 2d-FCS experiments. B.B. and G.T.D. contributed reagents. C.F. supervised the experiments. M.D.S.B., G.B., J.J.M., and A.S. wrote the manuscript.

