## Supplementary Information for "Apolipoprotein E4 has extensive conformational heterogeneity in lipid free and bound forms"

**corresponding author*

### Supplementary information.

### Methods.

#### Protein expression, purification, and labeling.

**Plasmid construct design.**

Apolipoprotein E protein (NCBI Reference Sequence: NP_000032.1 – for ε3 isoform) is expressed from a modified pET32a expression plasmid, which includes an N-terminal thioredoxin fusion protein, $His_6$ tag, and HRV 3C protease site followed by the Apolipoprotein E gene (excluding the first 18 residues, or signal peptide) as previously described[44]. The ApoE ε3 isoform gene was initially cloned into the BamHI EcoRI sites in the MCS of pET32a vector; however, the BamHI site was destroyed when the HRV 3C protease site was incorporated using site directed mutagenesis.

ATGAGCGATAAAATTATTCACCTGACTGACGACAGTTTTGACACGGATGTACTCAAAGCGGACGGGGCGATCCTCGTCGAT
TTCTGGGCAGAGTGGTGCGGTCCGTGCAAAATGATCGCCCCGATTCTGGATGAAATCGCTGACGAATATCAGGGCAAACTG
ACCGTTGCAAAACTGAACATCGATCAAAACCCTGGCACTGCGCCGAAATATGGCATCCGTGGTATCCCGACTCTGCTGCTG
TTCAAAAACGGTGAAGTGGCGGCAACCAAAGTGGGTGCACTGTCTAAAGGTCAGTTGAAAGAGTTCCTCGACGCTAACCTG
GCCGGTTCTGGTTCTGGCCATATGCACCATCATCATCATCATTCTTCTGGTCTGGTGCCACGCGGTTCTGGTATGAAAGAA
ACCGCTGCTGCTAAATTCGAACGCCAGCACATGGACAGCCCAGATCTGGGTACCGACGACGACGACCTGGAAGTGCTATTT
CAAGGACCCAAGGTTGAACAGGCTGTTGAAACTGAACCGGAACCCGAGCTGCGCCAGCAGACCGAGTGGCAGAGCGGCCAG
CGCTGGGAACTGGCACTGGGTCGCTTTTGGGATTACCTGCGCTGGGTGCAGACACTGTCTGAGCAGGTGCAGGAGGAGCTG
CTCAGCTCCCAGGTAACCCAGGAACTGAGGGCGCTGATGGACGAGACCATGAAGGAGTTGAAGGCCTACAAATCGGAACTG
GAGGAACAACTGACCCCGGTGGCGGAGGAGACGCGGGCACGGCTGTCCAAGGAGCTGCAGGCGGCGCAGGCCCGGCTGGGC
GCGGACATGGAGGACGTGCGCGGCCGCCTGGTGCAGTACCGCGGCGAGGTGCAGGCCATGCTCGGCCAGAGCACCGAGGAG
CTGCGGGTGCGCCTCGCCTCCCACCTGCGCAAGCTGCGTAAGCGGCTCCTCCGCGATGCCGATGACCTGCAGAAGCGCCTG
GCAGTGTACCAGGCCGGGGCCCGCGAGGGCGCCGAGCGCGGCCTCAGCGCCATCCGCGAGCGCCTGGGGCCCCTGGTGGAA
CAGGGCCGCGTGCGGGCCGCCACTGTGGGCTCCCTGGCCGGCCAGCCGCTACAGGAGCGGGCCCAGGCCTGGGGCGAGCGG
CTGCGCGCGCGGATGGAGGAGATGGGCAGCCGGACCCGCGACCGCCTGGACGAGGTGAAGGAGCAGGTGGCGGAGGTGCGC
GCCAAGCTGGAGGAGCAGGCCCAACAGATACGCCTGCAGGCCGAGGCCTTCCAGGCCCGCCTCAAGAGCTGGTTCGAGCCC
CTGGTGGAAGACATGCAGCGCCAGTGGGCCGGGCTGGTGGAGAAGGTGCAGGCTGCCGTGGGCACCAGCGCCGCCCCTGTG
CCCAGCGACAATCAC

MSDKIIHLTDDSFDTDVLKADGAILVDFWAEWCGPCKMIAPILDEIADEYQGKLTVAKLNIDQNPGTAPKYGIRGIPTLLL
FKNGEVAATKVGALSKGQLKEFLDANLAGSGSGHMHHHHHHSSGLVPRGSGMKETAAAKFERQHMDSPDLGTDDDDLEVLF
QGPKVEQAVETEPEPELRQQTEWQSGQRWELALGRFWDYLRWVQTLSEQVQEELLSSQVTQELRALMDETMKELKAYKSEL

EEQLTPVAEETRARLSKELQAAQARLGADMEDVRGRLVQYRGEVQAMLGQSTEELRVRLASHLRKLRKRLLRDADDLQKRL
AVYQAGAREGAERGLSAIRERLGPLVEQGRVRAATVGSLAGQPLQERAQAWGERLRARMEEMGSRTRDRLDEVKEQVAEVR
AKLEEQAQQIRLQAEAFQARLKSWFEPLVEDMQRQWAGLVEKVQAAVGTSAAPVPSDNH

Site-directed mutagenesis was performed on the pET32a Apolipoprotein E expression vector to create the ApoE (ε4 isoform) constructs (Supplementary Table 1). All mutations were verified using Sanger sequencing.

**Protein expression and purification.**

All ApoE4 constructs were expressed recombinantly in BL21-Gold (DE3) cells (Agilent). 2L cultures were grown in LB medium containing carbenicillin (100 µg/mL) to $OD_{600}$ ~0.8 and induced with 1 mM IPTG for 4 hours at 37 degrees C. Harvested cells were lysed with sonication at 4 degrees C in lysis buffer (50 mM Tris pH 8.0, 500 mM NaCl, 10 mg/mL lysozyme, 5 mM BME, and cOmplete™ EDTA-free Protease Inhibitor Cocktail (Roche)). The supernatant was cleared by centrifugation (37000 rpm for 1 hour) and loaded onto a HisTrap FF column (Cytiva) in buffer A (20 mM Sodium Phosphate pH 7.5, 500 mM NaCl, 20 mM Imidazole, 5 mM BME). The Thioredoxin-$His_6$-ApoE protein fusion was eluted with Buffer B (buffer A + 500 mM Imidazole) and dialyzed into HRV 3C protease cleavage buffer (50 mM HEPES pH 7.5, 100 mM NaCl, 5 mM BME) after adding HRV 3C protease to pooled fractions. Thus, the Thioredoxin-$His_6$ fusion protein was cleaved yielding full length ApoE4 with two additional N-term residues (GlyPro). FL ApoE4 was then bound to a Heparin Sepharose FF column (Cytiva) and eluted using a step gradient from 0 to 100% buffer B (buffer A: 50 mM HEPES pH 7.5, 100 mM NaCl, 5 mM BME, buffer B: buffer A + 1 M NaCl) over 140 min. Heparin elution fractions of FL ApoE4 were pooled and dialyzed overnight into buffer A (20 mM Sodium Phosphate pH 7.3, 2 M Urea, 5 mM BME) for subsequent ion exchange chromatography. Dialyzed FL ApoE4 was then bound to a Q Sepharose FF column (Cytiva) and eluted using a gradient of 0-40% buffer B (buffer A: (20 mM Sodium Phosphate pH 7.3, 2 M Urea, 5 mM BME, buffer B: buffer A + 1 M NaCl) over 80 min, then 40-100% buffer B over 60 min. Purified FL ApoE4 variants were analyzed using SDS-PAGE and verified by electrospray ionization mass spectrometry (LC-MS). Concentrations were determined spectroscopically in 50 mM Sodium Phosphate pH 7.5, 2 M Urea, 200 mM NaCl, 5 mM BME using an extinction coefficient = 44950 $M^{-1}$ $cm^{-1}$.

**Choice of labeling positions.**

The choice of labeling positions has been designed as a compromise between flanking regions of interest (hinge, C-terminal domain, four-helix bundle) and a series of different criteria regarding the structural properties of the protein. As there are no full-length structures available for either monomeric ApoE4 or ApoE4 oligomers, the ApoE3-like NMR structure (PDB: 2L7B) was used as a guide for identifying structural similarities with the resolved structure. In particular, we avoided altering amino acids clearly involved in structurally relevant interactions. When choosing labeling positions within folded domains, we focused on surface exposed residues to maximize accessibility of the cysteine residues during labeling. The spacing of the fluorophores was optimized to ensure use of the entire FRET dynamic range when the protein is unfolded, so that a contrast between the distance in the folded structure and in a disordered state would provide different transfer efficiencies. We also attempted to reduce the effects of quenching between fluorophores and aromatic residues[45,46]. With tryptophan residues identified as major quenchers for Alexa Fluor 488 and Alexa Fluor 594, fluorophores were positioned at least 10 residues away from its nearest tryptophan in the sequence.

**Protein labeling.**

All ApoE variants were labeled with Alexa Fluor 488 maleimide (Invitrogen Molecular Probes) under denaturing conditions in buffer A (20 mM Sodium Phosphate pH 7.3, 2 M Urea, 5 mM BME) at a dye/protein molar ratio of 0.7/1.0 at 4 degrees Celsius overnight. Single labeled ApoE protein was isolated using ion-exchange chromatography (Mono Q 5/50 GL, Cytiva – protein bound in buffer A and eluted with 0-40% buffer B (buffer A + 1 M NaCl) gradient over 80 min, then 40-100% buffer B over 20 min). UV-Vis spectroscopic analysis was used to identify fractions with 1:1 dye:protein labeling. Single labeled Alexa Fluor488 maleimide labeled N protein was then subsequently labeled with Alexa Fluor 594 maleimide at a dye/protein molar ratio of 1.2/1.0 overnight at 4 degrees Celsius. Double labeled (488:594) protein was then further purified using ion-exchange chromatography (Mono Q 5/50 GL, Cytiva –as listed above).

**Liposome preparation.**

Unilamellar vesicles were prepared from 1,2-dimyristoyl-sn-glycero-3-phosphocholine (DMPC) (Avanti Polar Lipids, USA) dissolved in Chloroform (25 mg/mL). The DMPC was used as received without further purification. DMPC solution containing 12.5 mg lipid was dried under a stream of dry nitrogen in glass scintillation vials. This lipid thin film was kept under vacuum overnight to remove traces of organic solvent. Liposomes were prepared by hydrating the film with 1 ml of 50 mM sodium phosphate buffer (pH 7.4) at 42 ± 1°C. The vial was shaken to form a rough liposomal suspension followed by five freeze-thaw cycles to obtain homogeneous preparation. Freezing was achieved by submersion into liquid nitrogen followed by thawing the sample in a water bath at 42 ± 1° C. The prepared vesicles were then extruded through an extruder set (Avanti Polar Lipids, USA) containing polycarbonate membranes with a pore size of 0.2 µm (Whatman plc., Cytiva Life Sciences). Finally, the extruded vesicles were flash frozen using liquid nitrogen and stored at -80°C for future use. Before the experiment, the frozen liposomes were thawed in a water bath at 42 ± 1°C followed by five freeze/thaw cycles, according to the protocol discussed earlier. The liposomes were then re-extruded using Avanti Polar Lipids extruder set at room temperature. The extruded liposomes were kept at 4°C and used within 2 days after the preparation. In parallel to single-molecule experiments, 50 µL of the prepared liposome solutions were used for cryo-TEM imaging using JEOL JEM-1400(Plus) (JEOL USA Inc., USA) 120 kV Transmission Electron Microscope. The images were captured using an AMT XR111 CCD camera and analyzed using a Graphical User Interface developed in Mathematica (Wolfram research).

### Data analysis.

**Experimental setup and procedure for single-molecule fluorescence experiments.**
Single-molecule fluorescence measurements were performed on a Picoquant MT200 instrument (Picoquant, Germany). For single-molecule FRET measurements, a Pulsed Interleaved Excitation (PIE) scheme was obtained by synchronizing a diode laser (LDH-D-C-485, PicoQuant, Germany) and a supercontinuum laser (SuperK Extreme, NKT Photonics, Denmark), filtered by a z582/15 band pass filter (Chroma) and pulsed at 20 MHz. Emitted photons were collected through a 60x1.2 UPlanSApo Superapochromat water immersion objective (Olympus, Japan). For nanosecond FRET-FCS measurements, the same diode laser was used in continuous-wave mode to excite the donor dye. Photons emitted from the sample were collected through a 60x1.2 UPlanSApo Superapochromat water immersion objective (Olympus, Japan), passed through a dichroich mirror (ZT568rpc, Chroma, USA), a long-pass filter (HQ500LP, Chroma Technology) to suppress scattering light. and a confocal pinhole (100 μm diameter). The emitted photons were then distributed into four channels, first by a polarizing beam splitter and then by a dichroic mirror (585DCXR, Chroma) for each polarization. Donor and acceptor emission was filtered (ET525/50m or HQ642/80m, respectively, Chroma Technology) and then focused on SPAD detectors (Excelitas, USA). The arrival time of every detected photon was recorded with a HydraHarp 400 TCSPC module (PicoQuant, Germany). FRET experiments were performed by exciting the donor dye with a laser power of 100 μW (measured at the back aperture of the objective). For PIE experiments, the power used for exciting the acceptor dye was adjusted to match a total emission intensity after acceptor excitation to the one observed upon donor excitation (between 50 and 70 μW). Single-molecule FRET efficiency histograms were acquired from samples with protein concentrations between 50 pM and 100 pM, estimated from dilutions of samples with known concentration based on absorbance measurements.
All measurements were performed in 50 mM NaPi pH 7.4, 143 mM β-mercaptoethanol (for photoprotection), 0.001% Tween 20 (for surface passivation) and GdmCl or TMAO at the reported concentrations. Tween 20 at this concentration is below the critical micelle concentration and we observe no evidence of interaction of Tween with the protein, neither interference with its ability of binding lipids. Measurements under aqueous buffer conditions are performed with PolyEthyleneGlycol (PEG)-passivated cuvettes[47], which significantly prevents adhesion of the protein to the surface. The exact concentration of denaturant is determined from measurement of the solution refractive index with an Abbe refractometer (Bausch & Lomb, USA).
Each sample was measured for at least 10 min at room temperature (295 ± 0.5 K).

**FRET efficiency histograms.**

Fluorescence bursts were identified by time-binning photons in bins of 1 ms and accepting bursts whose total number of photons after donor excitation was larger than at least 15 photons in each bin and contiguous bins were merged if the total number of photons was larger than at least 20 photons. The exact threshold was selected based on the background contribution identified in the photon counting histograms with 1 ms binning and a minimum common threshold across constructs has been used to minimize contributions to the width of transfer efficiency distributions due to different thresholds[30]. Transfer efficiencies for each burst were calculated according to

$$E = n_A/(n_A + n_D) \qquad \textbf{Eq. S1}$$

where $n_D$ and $n_A$ are the numbers of donor and acceptor photons, respectively.

Corrections for background, acceptor direct excitation, channel crosstalk, differences in detector efficiencies, and quantum yields of the dyes were applied[48]. The labeling stoichiometry ratio $S$ was computed accordingly to:

$$S = I_D/(\gamma_{PIE} I_A + I_D) \qquad \textbf{Eq. S2}$$

where $I_D$ and $I_A$ represent the total intensities observed after donor and acceptor excitation and $\gamma_{PIE}$ provides a correction factor to account for differences in the detection efficiency and laser intensities. Bursts with stoichiometry corresponding to 1:1 donor:acceptor labeling (in contrast to donor and acceptor only populations) were selected according to the criterion 0.3 < $S$ < 0.7. The final histogram of transfer efficiencies was constructed from these described selected bursts. Variations in the selection criteria for the stoichiometry ratio do not impact significantly the observed mean transfer efficiency (within experimental errors).

To estimate the mean transfer efficiency and extract multiple populations from the transfer efficiency histograms, each population was approximated with either a Gaussian or a LogNormal distribution function. For fitting more than one peak, the histogram was analyzed with a sum of the abovementioned functions. When analyzing multiple overlapping populations, in order to limit the model parameters and potential overfitting, we favored the use of global fit analysis where some parameters are shared across multiple or all concentrations. A list of used parameters is provided in the following.

For **ApoE4$_{182, 241}$**, data were globally fitted with the same width for each population across all concentrations of GdmCl. While this does not account for small deviations in the width due to the transfer efficiency dependence of the shot noise[30] , this option was made necessary because of the strong overlap between the two main populations across a large range of denaturant concentrations. The width was constrained between 0 and 0.09

based on the main width determined in high denaturant concentrations, where the region is completely unfolded. The upper limit value aligns with analogous values observed for other constructs.

For **ApoE4$_{223, 291}$**, we performed a global fit across all GdmCl concentrations with the following constraints: the mean transfer efficiency of the population at low transfer efficiency was shared across multiple GdmCl concentrations and constrained within a range of 0.06<E<0.18; the mean transfer efficiency of the population at high transfer efficiency was shared across multiple GdmCl concentrations and constrained within a range of E>0.85; the width of the distribution associated with each population was constrained for the population at intermediate transfer efficiencies between 0.06 and 0.13, whereas the one for the population at high transfer efficiency was constrained between 0 and 0.1.

For **ApoE4$_{86, 241}$**, no global fit was necessary but the mean transfer efficiencies of the three populations were constrained such that $0.1<E_1<0.25$, $0.3<E_2<0.75$, and $0.8<E_3<0.9$ for the low, intermediate, and high transfer efficiency population, respectively. All the widths were constrained between 0 and 0.12.

For **ApoE4$_{86, 165}$**, we initially fit the data using two Gaussian distributions and a Log-Normal distribution. After the initial fit suggests an almost constant value for the folded and intermediate population, the model has been further restrained to account for these fixed positions (though corrected for the refractive index change due to the increasing denaturant. Given the small amplitude, the width of the intermediate distribution has been enforced to be 0.1 across all denaturant concentration. Similarly, the width of the unfolded state has been constrained to 0.1 up to 2 M GdmCl, where the width of the distribution can start to be determined more robustly.

For **ApoE4$_{5,86}$**, we use a single Gaussian distribution without any constraint for each denaturant concentration above 1 M GdmCl. From 1 M GdmCl, we use a combination of a Log-Normal and a Gaussian distribution. Based on an iterative fitting procedure, further constraints have been enforced, requiring that $E_1 < 0.15$ and fixing the asymmetry of the Log-Normal distribution to 1.875. This is made necessary to limit the error in the estimate of the area of the low transfer efficiency population.

**Folding equilibrium.**

Folding equilibrium across the identified populations is described in terms of two-state and three-state models depending on the observable of the specific construct. Here, we exemplify the two-state model for the case of an unfolded (U) and folded (F) states and the three-state model as the equilibrium between native (N), intermediate (I) and unfolded (U) states. The same models can be applied to describe the folding equilibrium between different native states, as in the case of **ApoE4$_{223, 291}$**.

For the two-state equilibrium $\mathrm{N} \rightleftharpoons \mathrm{U}$, the corresponding fraction folded and unfolded can be written in terms of the equilibrium constant $K_{UN}$ as

$$f_U = \frac{1}{1+K_{UN}}, f_N = \frac{K_{UN}}{1+K_{UN}}$$ **Eq. S3**

where $K_{UN} = \exp[-\Delta G_0^{UN}/RT + m\,c] = \exp[m/RT(c - c_{1/2})] = \exp[\Delta G_0^{UN}/c_{1/2}(c - c_{1/2})]$, where $c$ is the denaturant concentration, $c_{1/2}$ represents the denaturant concentration at which the folding and unfolding fraction curves cross each other, e.g., when the two states have an equal abundance, $\Delta G_0^{UN}$ is the free energy difference between the N and U states extrapolated at zero denaturant concentration, R is the ideal gas constant and T is the temperature.

For the three-state equilibrium $\mathrm{N} \rightleftharpoons \mathrm{I} \rightleftharpoons \mathrm{U}$, the corresponding fraction folded, intermediate, and unfolded can be written in terms of the equilibrium constant $K_{UN}$ and $K_{NI}$ as

$$f_U = \frac{1}{1+K_{UI}+K_{UI}\,K_{IN}}, f_I = \frac{K_{UI}}{1+K_{UI}+K_{UI}\,K_{IN}}, f_N = \frac{K_{UI}\,K_{IN}}{1+K_{UI}+K_{UI}\,K_{IN}}$$ **Eq. S4**

where $K_{UI} = \exp[\Delta G_0^{UI}/c_{1/2}^{UI}(c - c_{1/2}^{UI})]$ and $K_{IN} = \exp[\Delta G_0^{IN}/c_{1/2}^{IN}(c - c_{1/2}^{IN})]$ with $\Delta G_0^{UI}$ and $\Delta G_0^{IN}$ represent the free energy difference extrapolated to aqueous buffer conditions between the U and I and I and N states, respectively, whereas $c_{1/2}^{UI}$ and $c_{1/2}^{IN}$ are the concentration where the corresponding fraction curves cross each other. In this case, it is important to notice that the crossing point does not necessarily occur in the midpoint (50%) of the transition.

Finally, when the folding transition is not reported in terms of fractions but is estimated based on a specific signal (e.g., variation in mean transfer efficiency or width of the transfer efficiency distribution), the total signal is described as combination of specific fraction with the corresponding signal of each population, where the signals associated to each population are considered further parameters.

**IRF determination.** Instrument response function (IRF) for each 4 detectors and for the 2 lasers was obtained recording the TCSPC histogram for the fluorescence of Rhodamine B (RhB, 100 μM) in presence of high concentration of potassium ferricyanide (95% of a saturated solution of $K_3Fe(CN)_6$ at 23°C in water), an electron-transfer fluorescence quencher[49].

As the rate of non-radiative relaxation pathways of the fluorophore increases with addition of quencher, the overall fluorescence time decay gets faster and eventually it is fast enough in comparison with the IRF -effectively a delta function-, the observed TCSPC histogram approaches the actual IRF[50]. This is due to the fact the observed TCSPC histogram is a time convolution of the IRF and the fluorescence relaxation kinetics of the fluorophore,

$$f(t) = \int_0^t IRF(t-\tau)I(\tau)d\tau \quad \textbf{Eq. S5}$$

and the convolution of a function with a δ-function is equal to the function itself.

Laser power was set at 0.5 μW after the main dichroic at a repetition rate of 20 MHz and decreased 100 times before measurement (final power ~ 5 nW). Data was collected with 16 ps resolution. The collected TCSPC histograms were corrected by subtracting dark counts - which appear uniformly distributed in the histogram - and normalized over the total number of counts.

For consistency, we checked if at the ferricyanide concentration used its quenching effect on the TCSPC histogram was already maximum in the conditions of our measurements. On the one hand, we found that the histograms converged and became sharper as the concentration was increased and from 0.8 M up, they were indistinguishable. On the other hand, we also found that, as the mean fluorescence intensity decreased, the computed steady-state anisotropy converged to a value of 0.37-coincident with previously reported values of limiting anisotropy of RhB derivatives [51], as expected for emission taking place much faster than rotational relaxation from randomly oriented fluorophores with almost parallel excitation and emission transition dipolar moments[52].

We found stronger quenching and faster fluorescence decay of RhB with ferricyanide than with potassium iodide, a quencher commonly used for this purpose, both tested at the concentration of a 95% saturated solution at room temperature (~23°C). Additionally, in presence of quencher, we found no detectable differences in the normalized TSCPC histograms obtained with Alexa594 upon 589 nm excitation with 640 nm detection, but slightly slower fluorescence decay with Alexa488 in comparison to RhB, upon 485 nm excitation and 525 nm detection.

**Correction factors for quantitative anisotropy determination.** In order to obtain estimates of the correction coefficients $L_p$ and $L_s$ accounting for scrambling of polarization due to tight focusing optics we employed the technique developed by Koshioka and coworkers[53]. The methodology consists in performing a measurement of fluorescence intensity in two solutions of a fluorophore, in low and high viscosity media, recording the signal in 2 detectors collecting light polarized parallel (p) and perpendicular (s) to that of the excitation laser. The time correlated single photon counting (TCSPC) histogram of p- and s- polarization detection for each medium is fitted as a convolution with the Instrument Response Function (IRF) and a fluorescence lifetime function containing fluorescence and rotational relaxation kinetic terms. The parameters $L_p$ and $L_s$ are obtained through global fitting of the TCSPC histograms from the two solutions and two polarizations.

To this end, we used maleimide-C5 Alexa488 to determine $L_p$ and $L_s$ in detectors for donor dye upon excitation at 485 nm, and maleimide-C5 Alexa594 dyes to determine $L_p$ and $L_s$ in acceptor-channel detectorsupon excitation at 590 nm. The concentration of dye solutions was 5 μM in water for the low viscosity solution and in

59% v/v glycerol in water (high viscosity solution), supplemented with 240 mM beta-mercaptoethanol. Measurements were performed at 23°C ± 1°C. Laser power was set at 0.5 μW after the main dichroic at a repetition rate of 20 MHz and decreased 100 times before measurement (final power ~ 5 nW). Data was collected with 16 ps resolution.

The following equations for parallel (*p*) and perpendicular (*s*) detection were simultaneously fit to the corresponding pair of experimental TCSPC histograms:

$$f_p^d(t) = \Delta\tau \sum_{j=1(\tau_j=0)}^{k(\tau_k=t)} IRF_p^{d,\lambda}(\tau_j - t)\; I_p(\tau_j)$$
$$f_s^d(t) = \Delta\tau \sum_{j=1(\tau_j=0)}^{k(\tau_k=t)} IRF_s^{d,\lambda}(\tau_j - t)\; I_s(\tau_j)$$ **Eq. S6**

where *d* denotes the detector, *λ* denotes the excitation laser employed, $IRF^{d,\lambda}$ is the corresponding experimentally determined instrument response function, and $I_p$ and $I_s$ are given by,

$$I_p(\tau) = a_p\, e^{-\frac{\tau}{\tau_F}} \left[1 + r_0\left(2 - 3L_p\right) e^{-\frac{\tau}{\tau_R}}\right]$$
$$I_s(\tau) = a_s\, e^{-\frac{\tau}{\tau_F}} \left[1 - r_0\left(1 - 3L_s\right) e^{-\frac{\tau}{\tau_R}}\right]$$ **Eq. S7**

where $a_p$ and $a_s$ are amplitude coefficients, $\tau_F$ is the fluorescence emission lifetime parameter, $\tau_R$ is a rotational diffusion characteristic time parameter, $r_0$ is the intrinsic maximum fluorescence anisotropy parameter, and $L_p$ and $L_s$ are the corresponding scrambling correction coefficients for each polarization. Parameters $a_p$, $a_s$, $\tau_F$, $\tau_R$, $L_p$ and $L_s$ were floating fitting parameters while $r_0$ was fix at the previously reported value for Alexa488 and Alexa594, ~ 0.38 [54]. Simultaneous fitting of **Eq. S7** to the pairs of TCSPC histograms obtained in water and in 59% v/v glycerol was performed sharing only parameters $L_p$ and $L_s$, employing weighted least squares, using 1/*f*(t) as weights.
Computing the time resolved anisotropy, *r*(t),

$$r(t) = \frac{F_p(t) - G\,F_s(t)}{\left(1 - 3L_s\right)F_p(t) + \left(2 - 3L_p\right) G\,F_s(t)}$$ **Eq. S8**

also requires estimation of the relative efficiencies of detection in both detectors, parallel and perpendicular, denoted by the *G* factor, i.e., the efficiency of detection compared between parallel and perpendicular polarization.

To this end, for each pair of detectors, we used 3 different approaches which show consistency with each other:
- the ratio of amplitude parameters $a_p$ and $a_s$ in **Eq. S7**;
- solving **Eq. S8** for $G$ using independent measurements of steady-state anisotropy together with measurements in our microscope of time-average fluorescence intensity $<F_p>$ and $<F_s>$ and parameters $L_p$ and $L_s$ obtained as described above.
- fitting the value of $G$ such that the time trace of $r(t)$ in **Eq. S8** obtained for free Alexa dyes in water at room temperature, decreases asymptotically to 0 and effectively reaches that value in less than 10 ns.

The values obtained of $L_p$, $L_s$ and $G$ for each excitation laser and each pair of detectors and employed in the analysis of the results in this work are shown in **Supplementary Table 5**.

**Fluorescence lifetimes analysis.** We estimated the fluorescence lifetime of the donor and acceptor by globally fitting the histogram of photon arrival times with **Eq. S7**, where $a_p$ and $a_s$ are treated as free parameters, $c_0$ is fixed to 0.38 as an average value based on previous estimates for Alexa Fluor 488 and Alexa Fluor 594[54] and $\tau_F$ and $\tau_r$ are the lifetime of the fluorophore and the rotational component of the dye, respectively. To this end, we time-gate the fluorescence arrival time to separate photons derived from donor and acceptor excitation and we select for stoichiometry and transfer efficiency to restrict the analysis to each subpopulation. We found that addition of an additional fitting parameter representing background counts is particularly useful when analyzing high transfer efficiency populations, such as for **ApoE$_{86,165}$**. Obtained values are reported in **Supplementary Table 6** and **7** for lipid-free and lipid-bound constructs. The lifetime of donor in presence or absence of acceptor is used to further compute the FRET rate. This provides an overall measurement of the characteristic timescales at play, enabling to compare whether the energy transfer occurs on timescale longer than the rotation of the fluorophores, which is commonly assumed to interpret transfer efficiency according to:

$$E = \int_{b_0}^{l_c} E(r)P(r)dr \qquad \textbf{Eq. S9}$$

Where $b_0$ is the shortest contact distance and $l_c$ is the contour length of the chain assuming all amino acids are in part of a disordered polymer (i.e., equal to the number of amino acids times the $C_\alpha$-$C_\alpha$ distance).

**Anisotropy analysis.** Anisotropy of the donor-only, acceptor-only, and acceptor after donor excitations are computed based on **Eq. S8** from photon arrival times histograms detected on the *p*- and *s*- polarization. Time-resolved anisotropy decays are then fitted to a double exponential decay according to:

$$\left((c_0 - c_\infty)\, e^{-\frac{t}{\tau_r}} + c_\infty\right) e^{-\frac{t}{\tau_M}} \quad \textbf{Eq. S10}$$

where $c_0$ is fixed to 0.38 as described above, $c_\infty$ is the residual anisotropy at infinite time, $\tau_r$ represents the rotational component of the dye, and $\tau_M$ reports about tumbling of the overall molecule. for the starting time distribution to account for the contribution. Paralleling the analysis of lifetime fluorescence, values for $c_\infty$ and $\tau_r$ are reported in **Supplementary Tables 8** and **9.** $\tau_r$ can be compared to the analogous quantity obtained from lifetime fits and is always faster than the corresponding inverse of the FRET rate or donor lifetime. $c_\infty$ are used to compute the orientation factor $\kappa^2$. Finally, we estimated also steady-state anisotropies by computing anisotropy based on the burst-determined photons in the different polarizations. The final calculation is equivalent to the one presented in **Eq. S8**, where the time-dependent factors are replaced by the number of photons in each selected burst.

***Orientation Factor $\kappa^2$*.** While commonly assumed to be equal to 2/3, the orientation factor $\kappa^2$ can be estimated using residual fluorescence anisotropies, which provide boundaries on the angles sampled by fluorophores. Though this is subject to various approximations, it does provide a useful test to quantify whether the accessible space sampled by the dyes is hindered by a folded domain. Following previous treatments where tumbling of the dye is described as a wobble-in-a-cone [54,55], $\kappa^2$ can be estimate as:

$$\kappa^2 = \left(1 - \sqrt{\frac{r_A}{r_0}}\right)\left(\sqrt{\frac{r_D}{r_0}}\left(\cos\frac{\theta_+ - \theta_-}{2}\right)^2 + \frac{1}{3}\right) +$$

$$\left(1 - \sqrt{\frac{r_D}{r_0}}\right)\left(\sqrt{\frac{r_A}{r_0}}\left(\cos\frac{\theta_+ + \theta_-}{2}\right)^2 + \frac{1}{3}\right) + \frac{1}{r_0}\sqrt{r_A r_D}\left(\frac{3}{2}(\cos\theta_+ + \cos\theta_-) - \cos\beta\right)$$

**Eq. S11**

where $\cos\beta = \sqrt{\frac{2}{3}\frac{r_{A(D)}}{r_A r_D} + \frac{1}{3}}$. Based on previous determinations, $r_0$ is set to 0.38. The only two unknowns are the angles $\theta_+$ and $\theta_-$, whose boundaries are estimated as:

$$\cos^{-1}(\min(\cos\beta, 1)) < \theta_+ < 2\pi - \cos^{-1}(\min(\cos\beta, 1)) \quad \textbf{Eq. S12a}$$

$$-\cos^{-1}(\min(\cos\beta, 1)) < \theta_- < \cos^{-1}(\min(\cos\beta, 1)) \quad \textbf{Eq. S12b}$$

with $\min$ indicating the smaller among the two values. Evaluating the solution across all the possible angles in these two intervals enables reconstructing a distribution of the possible $\kappa^2$ values, which deviates from the expected distribution for freely rotating fluorophores. Given that the tumbling of fluorophores occurs on a time scale faster the fluorophore lifetime and that of the inverse of the rate of energy transfer (see **Supplementary Table 6-9**), we can compute a mean value of $\kappa^2$ from the distribution, which can be compared to the expected value of 2/3. All the measured values are within the range of 0.74-1.01, with a bias toward higher values (stronger hindrance) for the lipid-bound states. It is important to note that $\kappa^2$ enters in the Forster radius at the power of 1/6 and therefore such deviations from 2/3 have an impact of about 5% on the estimate value of $R_0$ (**Supplementary Table 10-11**).

From the distributions of $\kappa^2$, we can further evaluate the precision and accuracy associated to the estimate of mean $\kappa^2$, as well as the minimum and maximum of the distribution:

$$\text{Precision} = \int_{k_{min}^2}^{k_{max}^2} dk^2 \left(1 - \left(\frac{3}{2}k^2\right)^{-1/6}\right) P(k^2) \Big/ \int_{k_{min}^2}^{k_{max}^2} dk^2 P(k^2) \qquad \textbf{Eq. S13}$$

$$\text{Accuracy} = \text{var}\left(\left(\frac{3}{2}k^2\right)^{-1/6}\right) \qquad \textbf{Eq. S14}$$

$$k_{min}^2 = \frac{2}{3}\left(1 - \left(\sqrt{\frac{r_D}{r_0}} + \sqrt{\frac{r_A}{r_0}}\right)/2\right) \qquad \textbf{Eq. S15}$$

$$k_{max}^2 = \frac{2}{3}\left(1 + \sqrt{\frac{r_D}{r_0}} + \sqrt{\frac{r_A}{r_0}} + 3\sqrt{\frac{r_D}{r_0}}\sqrt{\frac{r_A}{r_0}}\right) \qquad \textbf{Eq. S16}$$

The minimum and maximum of $\kappa^2$ are analytically computed assuming there is no knowledge of $r_{\infty-A(D)}$. A computational value can be obtained by evaluating all the possible angles allowed by **Eq. S12**.

The terms $r_A, r_D, r_{A(D)}$ reflects the residual anisotropy of the acceptor only, donor only and acceptor after donor excitation, respectively, and can be estimated from the values of residual anisotropy associated with the dye relaxation $r_{\infty-Aonly}, r_{\infty-Donly}, r_{\infty-A(D)}$ (see previous section). As validation, we further compared the values obtained with this method with analogous estimates obtained from the corresponding steady-state anisotropies, which is often used as a first approximation for the estimate of the residual anisotropies. Importantly, steady-state anisotropies are computed as an average across population specific bursts (donor-only, acceptor-only, donor-acceptor) and therefore provide an independent estimate compared to the time resolved ones. We found a general good agreement between these two methods of estimate, suggesting that the quantification of anisotropies and $\kappa^2$ are robust.

For the conversion of transfer efficiency to distances, we then use the value of the Förster radius for Alexa Fluor 488 and Alexa Fluor 594 previously determined and reported in literature, $R_0$ = 5.4 nm[56], corrected by the variation in solution refractive index and $\kappa^2$.

**Lifetime vs FRET.** The fluorescence lifetime is related to the mean transfer efficiency[30] through:

$$\tau_{DA}/\tau_D = 1 - \langle E \rangle + \frac{\sigma^2}{1-\langle E \rangle} \quad \textbf{Eq. S17}$$

where $\sigma^2$ is related to the variance of the sampled distribution *via*:

$$\sigma^2 = \int_0^\infty E(r)^2 P(r) dr - \langle E \rangle^2 \quad \textbf{Eq. S18}$$

For $\sigma^2$ equal to zero, Eq. S17 reduces to the linear trend expected for a rigid distance. For fitting experimental data, we choose the $P(r)$ of a wormlike chain[57], as described by:

$$P_{WLC}(r, l_p, l_c) = \frac{4\pi (r/l_c)^2 C(l_p, l_c)}{l_c(1-(r/l_c)^2)^{9/2}} \mathrm{Exp}\left[-\frac{3\, l_c}{4 l_p (1-(r/l_c)^2)}\right] \quad \textbf{Eq. S19a}$$

$$C(l_p, l_c) = \frac{1}{\pi^{3/2} e^{-\alpha} \alpha^{-3/2} (1 + 3/\alpha + 15/(4\alpha^2))} \quad \textbf{Eq. S19b}$$

with $\alpha = \frac{3}{4}\frac{l_c}{l_p}$, where $l_p$ is the persistence length. In analyzing the lifetime vs transfer efficiency data, both $l_p$ and $l_c$ are varied. Whereas for a given polymer the contour length is fixed, using the contour length as a fitting parameter for a region in which some parts are folding reflects the inherent change in the nature of polymer: the polymer cannot be anymore stretched to the say contour length because of the folded regions. It is an approximation used to provide an empirical quantification of the underlying distance distribution as well as to provide a qualitative measurement that accounts for the conformational changes within the protein. The limit of a completely dynamic chain is recovered for a Gaussian distribution:

$$P_G(r, \langle r^2 \rangle) = 4\pi r^2 \left(\frac{3}{2\pi \langle r^2 \rangle}\right)^{3/2} \mathrm{Exp}\left[-\frac{3}{2}\frac{r^2}{\langle r^2 \rangle}\right] \quad \textbf{Eq. S20}$$

**MD simulations of lipid-free ApoE4.** The NMR structure of ApoE3 (2L7B) was used as a starting model for all our simulations. Using pymol's mutagenesis wizard, we restored the native, mature, sequence of ApoE3 and made subsequent mutations from this model to acquire structures for ApoE2 (R158C), ApoE4 (C112R), and ApoE3 Christchurch (R136S). Each model was solvated in a dodecahedron box with 2.1 nm between the protein and the edge of the box. Systems were subsequently solvated, $Na^+$ and $Cl^-$ ions were added to a final concentration of 0.1 M with no net system charge, and the system was energy minimized with a steepest descents algorithm until the maximum force fell below 100 kJ $mol^{-1}$ $nm^{-1}$, using a step size of 0.01 nm and a cut-off distance of 1.2 nm for the Coulomb interactions, van der Waals interactions, and neighbors list. The AMBER03 force field[58] and explicit TIP3P solvent[59] were used for all simulations. Systems were subsequently equilibrated for 1.0 ns, where all bonds were constrained with the LINCS algorithm[60] and virtual sites to allow for a 4.0 fs timestep. Cut-offs of 1.1 nm were used for the neighbor list with 0.9 for Coulomb and van der Waals interactions. We deployed the particle mesh Ewald method for treatment of long-range interactions with a Fourier spacing of 0.12 nm. The Verlet cut-off scheme was used for the neighbor list. The stochastic velocity rescaling (v-rescale) thermostat[61] was used to hold the temperature at 300K. All simulations were prepared using Gromacs[62] 2020.

We ran our FAST adaptive sampling algorithm[43] on each of these models to explore distances between the following residue pairs: R92-W264, P183-K242, and R224-A292. The FAST algorithm balances directed exploration and unbiased simulations to efficiently explore conformational space of proteins. The algorithm proceeds as follows: 1) run initial simulations, 2) build a Markov State Model (MSM) for the aggregate simulation time, 3) rank each observed state in the MSM based on the exploration parameter, 4) restart simulations from the top ranked states, 5) repeat steps 2-4 until the specified number of FAST rounds is complete. For each model, we ran our FAST-sampling algorithm at 300 K for 12 rounds, followed by another 8 rounds of FAST-string (20 total) with 10 simulations per round[63]. Each simulation was 40 ns in length, for a total simulation time of 8 μs for each variant. For ApoE4 we performed 2 independent rounds of the above FAST sampling. All FAST simulations were performed using Gromacs.

As ApoE is highly flexible, we wanted to ensure we had robust sampling to adequately calculate the probability of each state. Using ENSPARA's clustering app[64], we clustered the FAST simulations from all mutants into a shared state space model coarse-grained to a final RMSD of 3.5 Å. This clustering resulted in a total of 18,182 discrete states. As each cluster center could come from any ApoE variant, we utilized MODELLER[65] to mutate all sequences back to ApoE4. Subsequently, we solvated each state in a dodecahedron box whose edges extend 1.0 nm beyond ApoE. The remainder of the system preparation followed that of the FAST system preparation. We next launched five independent simulations from each structure on the distributed computing platform, Folding@home. Each trajectory ran for a maximum of 100 ns, though the average trajectory length was 37 ns, for an aggregate simulation time of 3.43 ms. Simulations were performed using OpenMM[66], using the Langevin Integrator, a step size of 2 fs, and a temperature of 300K. During our simulations, some trajectories resulted in protein unfolding and caused ApoE to interact with its periodic image. These trajectories, along with trajectories that had improper periodic boundary condition removal, were excluded from our analysis resulting in a total of 3.37 ms of aggregate simulation time analyzed.

As the C-terminal domain of ApoE is highly flexible, we were concerned that clustering on all-atom features such as RMSD would yield a poorly connected state-space. Accordingly, we clustered ApoE on the distances between 15 different residue pairs distributed throughout ApoE. We selected these pairs to both ensure that we had an overall understanding of various aspects of ApoE movement, including extension of the C-terminal domain, movement of the N-terminal domain, motions between each domain, and the five distances measured in our single-molecule FRET experiments. The total list of residue pairs is in **Supplementary Table 16**. Following coarse-graining, we generated a Markov State Model (MSM) using Enspara's MSMBuilder[64] with a lag time of 10 ns.

**Post-hoc calculation of FRET histograms.** FRET histograms were calculated through a kinetic Monte Carlo simulation using the constructed Markov State Model (MSM). We designed this simulation to replicate the experimental process of single molecule FRET. First, we model dyes onto each structure in the MSM, removing any dye positions that touch or overlap with the protein, and finally compute the distribution of distances between each potential dye position.

Next, we simulate smFRET by recoloring an experimental photon burst. To do this, we select the photon arrival times from an experimental FRET trace and simulate a synthetic trajectory from our MSM of length αt, where α is a time rescaling factor and t is the total length of time of the experimental FRET burst. The states visited during the synthetic trajectory are governed by the transition probabilities of our MSM model. At each photon arrival time in the experimental burst, we select the corresponding state in our synthetic trajectory and randomly choose a position of the FRET dyes based on the previously calculated distribution probability for that state. To determine whether the observed photon is the result of a FRET event, or a donor emission, we convert the inter-dye distance into a FRET transfer probability based on the following equation:

$$E(r) = R_0^6/(R_0^6 + r^6) \qquad \textbf{Eq. S21}$$

In each case, the transfer probability is compared to a random number between 0 and 1 to determine if the photon was emitted as a donor or acceptor photon. This process is repeated for each photon event in the trajectory, and the resulting acceptor photons are summed and divided by the total number of photons to yield the FRET efficiency for the simulated burst. This process is repeated for all bursts in our experimental photon trace and the resulting FRET efficiencies are summed and plotted as the displayed FRET histograms. As smFRET is highly sensitive to changes in timescale and MD simulations can be faster than experimental time, we fit the time rescaling factor, α, using the experimental FRET distributions (see **Supplementary Figure 16).** The code for this kinetic Monte Carlo simulation has been developed as a command line app and is distributed via the enspara github (https://github.com/bowman-lab/enspara).

**Coarse-grained model of lipid-bound states.** While an atomically-detailed description of the protein requires all-atom simulations that are beyond the scope of this work, to provide a possible interpretation of the lipid-bound states observed with FRET measurements, we performed high coarse-grained MonteCarlo simulations of the protein that satisfy the FRET constraints. The scope of these simulations is purely illustrative and does not account for specific interactions with lipids or within the protein; instead, we aim only to provide a possible visual interpretation of the FRET data. To this end, we simplify the structure of the protein to a set of random coils (corresponding to the disordered regions) and rigid sticks (corresponding to the alpha helices), under the assumption that these elements are maintained in the lipid-bound state. Random coils are simulated by identifying the length of the segment from a distribution that follows the statistics of:

$$P(r,N) = (1/0.18\,N^{-3/2})4\pi r^2 \mathrm{Exp}[-3/2\,r^2/(0.38 * 0.4 * N)] \qquad \textbf{Eq. S22}$$

based on previous estimates for a disordered region[67]. Since disordered segments are often significantly shorts, their main effect is to contribute to protein flexibility and, therefore, we neglect effects of local compaction or expansion of disordered regions due to differences in sequence composition. To enforce bound geometries, we reject all configurations that are outside a spherical corona of radius 8.67 nm and thickness 0.1 nm. For disordered regions this entails checking whether the beginning or end of the segment are within the spherical corona. For the rigid regions, this is performed for the end, middle point, and beginning of the rigid region. We further allow for overlapping configurations of the protein to facilitate the search of configurations in which helices could come in contact. We start the simulation by simulating the protein region between 86 and 241, since it displays heterogeneity in the experimental data and entails three experimental constraints (as given by **ApoE4$_{86,165}$**, **ApoE4$_{182,241}$**, and **ApoE4$_{86,241}$**). Out of 500,000 productive results of the simulation, only 1 configuration is found to satisfy the experimental results for the expanded state of **ApoE4$_{86,165}$** and only 2 configurations satisfy the other state. We then append to the resulting states an additional 50,000 configurations of the residues between 1 and 86 and 50’000 configurations of the residues between 241 and 299. By using the additional experimental constraints, we obtain a total of 39249 configurations of the protein, where 4361 elements differ for the N-terminal tail and 9 for the C-terminal domain. The lowest number of configurations identified in the C-terminal domain stems from the larger number of constraints imposed on position 241. Two examples of configurations are reported in **Fig. 5** to provide an interpretative model of the lipid-bound state of the protein.

### Additional notes.

#### ApoE oligomerization.

Previous work on oligomerization kinetics of ApoE using ensemble FRET has provided estimates for the dimerization and tetramerization rates of ApoE4 and corresponding equilibrium dissociation constants for dimer and tetramer[18]. The dissociation constant between the monomer and dimer state is determined to be $K_D^{mon-dim}$ ~ 80 nM, whereas the dissociation constant between dimers and tetramers is $K_D^{dim-tet}$ ~ 20 nM. As a result, previous analysis (compare Fig. 8d in [18]) suggests that:

- at 10 nM of total protein concentration, ~20% of ApoE4 molecules form dimers;
- at 100 nM of total protein concentration, ~30% of ApoE4 molecules form dimers and an additional ~30% is assembled in tetramers;
- at 1 μM of total protein concentration, ~80% and ~15% of ApoE4 molecules are in the tetramer and dimer forms, respectively;
- at 10 μM of total protein concentration, ~95% of ApoE4 molecules are part of tetramer species.

### Supplementary Figures.

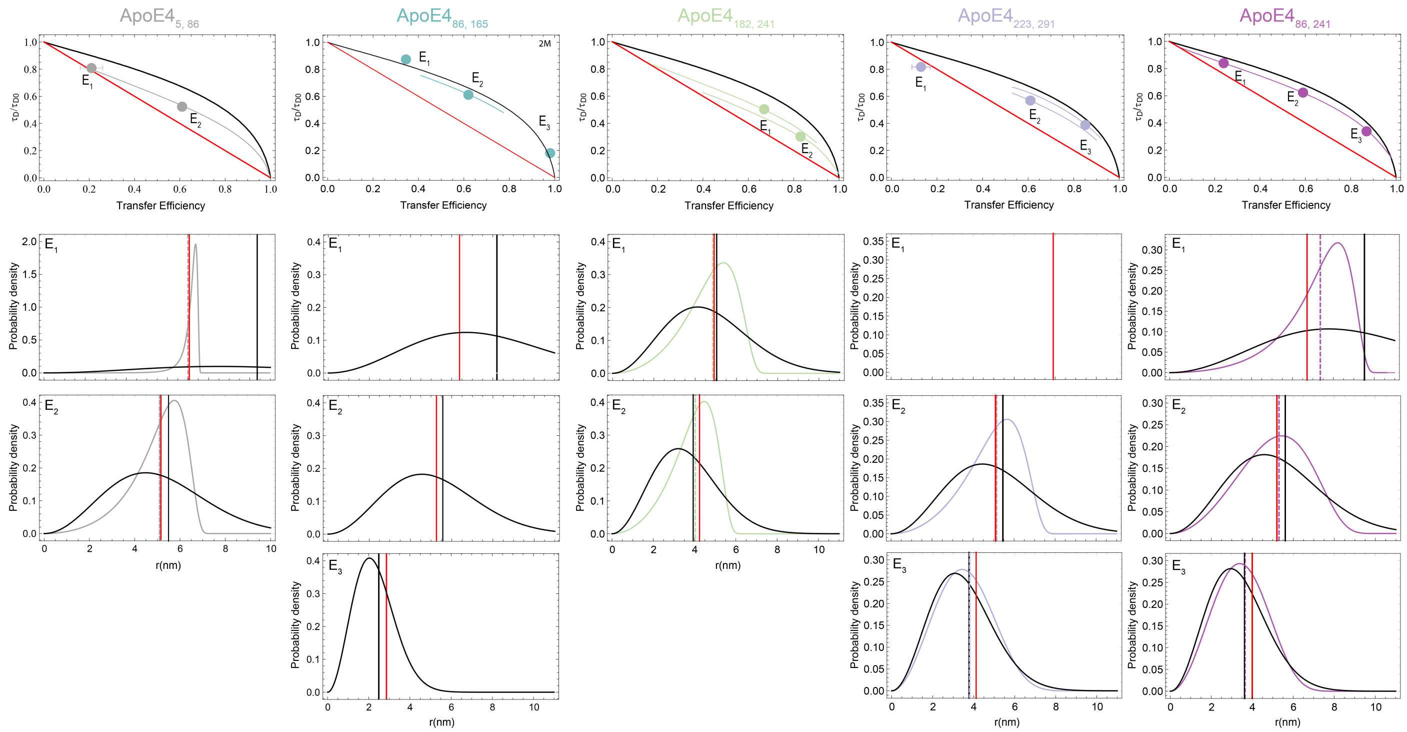


**Supplementary Figure 1**: **Lifetime *vs* transfer efficiency plot and corresponding distance distributions for each of the five ApoE4 constructs measured in aqueous buffer conditions (**50 mM NaPi pH 7.4). The solid red line indicates the expected result for a rigid distance. The solid black line reports the expected trend for a Gaussian chain distribution whose mean square end-to-end distance satisfies the measured mean transfer efficiency. The colored line represents the best fit to a wormlike chain, where both persistence length and contour length are fitted to satisfy the constraints imposed by the measured mean transfer efficiency and mean lifetime.

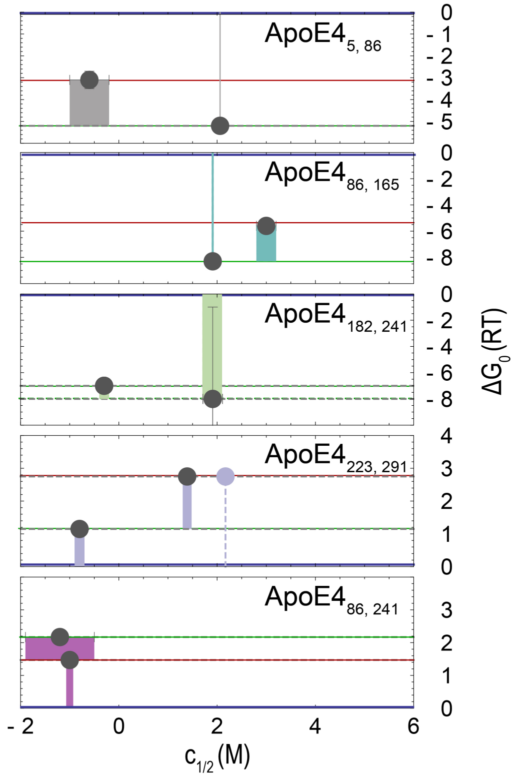


**Supplementary Figure 2. Folding free energy and midpoint of the transition determined from the folding equilibrium analysis.** Plot of fitting parameters in **Supplementary Table 2** obtained from the folding equilibrium analysis of **Fig. 2** with **Eq. S4**. Blue, red, and green line represent the corresponding states from which the $\Delta G_0$ are computed. Gray (**ApoE4$_{5,86}$**), teal (**ApoE4$_{86,165}$**), green (**ApoE4$_{182,241}$**), purple (**ApoE4$_{223,291}$**) and violet bars (**ApoE4$_{86,241}$**) connect the states associated with the computed $\Delta G_0$. Width of the bars represents the associated error on the corresponding $c_{1/2}$, whereas vertical error bars quantify the error (from the fit) associated with $\Delta G_0$. Dashed vertical line in **ApoE4$_{223,291}$** identifies the transition in **Supplementary Fig. 4** when analyzed with a two-state model.

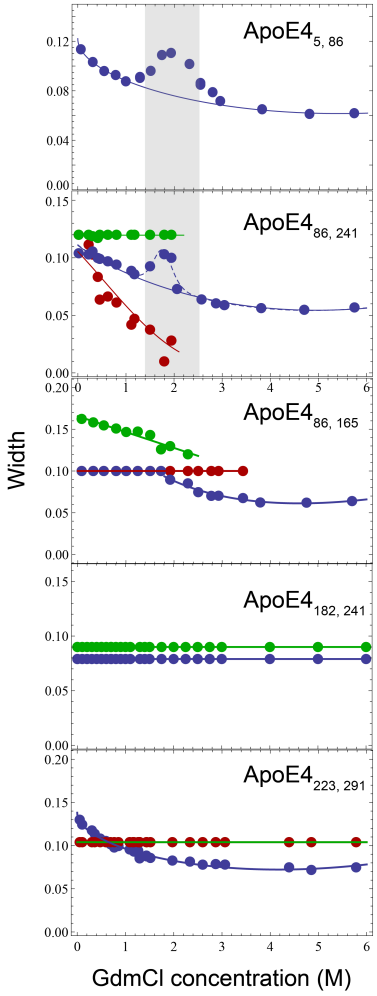


**Supplementary Figure 3**: Widths of transfer efficiency distributions attributed to independent populations in each construct. Solid lines are polynomial fits to help guiding the eyes. The widths of **ApoE4$_{5,86}$** and **ApoE4$_{86,241}$** exhibit an increase between 1.25 and 2.5 M GdmCl suggesting coexistence of two independent configurations under the same peak, which we interpret as a conformational shift occurring during the folding of the four-helix bundle. In **ApoE4$_{182,241}$** the widths of the two populations are best described using a shared parameter for each population in the global fit of the denaturant titration. In **ApoE4$_{223,291}$**, the width of the main population increases when decreasing GdmCl concentration from 1 M to 0 M, which is compatible with the range where previous experiments identified formation of structure in the C-terminal domain. Due to the small amplitudes of the low and high transfer efficiencies populations in **ApoE4$_{223,291}$**, the corresponding widths were fixed across all concentrations. Similarly, in **ApoE4$_{86,165}$**, due to the small amplitude of the unfolded and intermediate state, below 3 M GdmCl the widths of the corresponding transfer efficiency distribution were fixed. Errors associated with repeated measurements are reported in **Supplementary Table 1**.

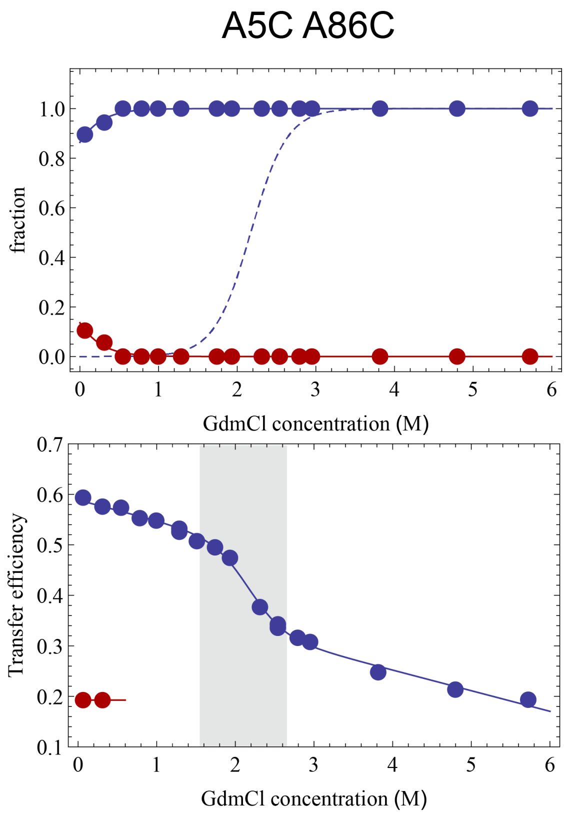


**Supplementary Figure 4**. Analysis of the **ApoE4$_{5,86}$** construct using one single population in the range from 0.5 and 6 M GdmCl. When fitting the transfer efficiency with one single population, we observe a jump in transfer efficiencies (lower panel) and a broadening of the width (see **Supplementary Fig. 3**) in the range between 1.5 and 2.5 M GdmCl. The range of the transition corresponds to the folding transition of the four-helix bundle. Corresponding values of the fit are reported in **Supplementary Table 2**.

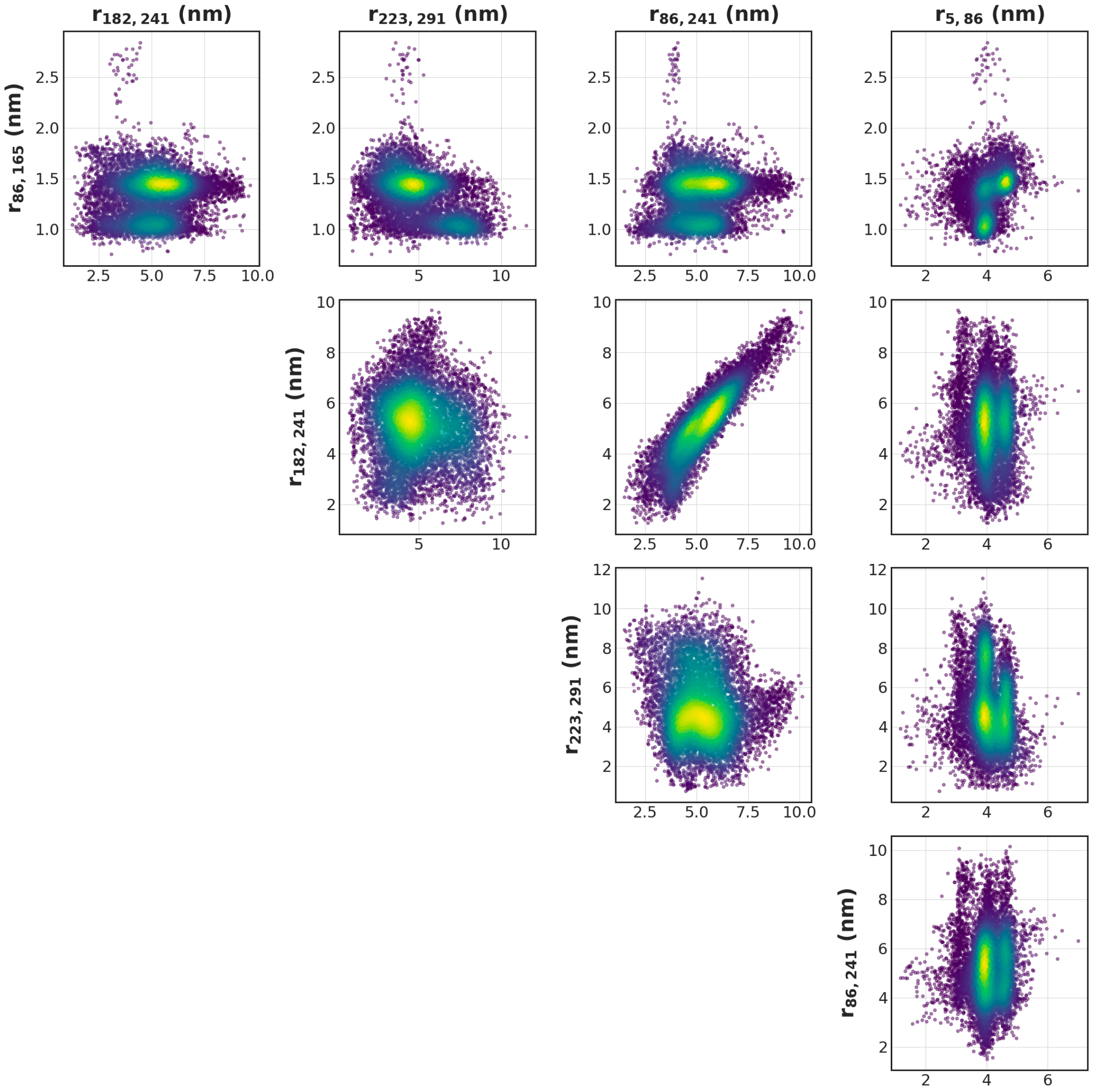


**Supplementary Figure 5. Correlations across different pair distances in lipid-free ApoE4. .** Distance pair correlations from MD simulations contrasting all labeled distance pairs.

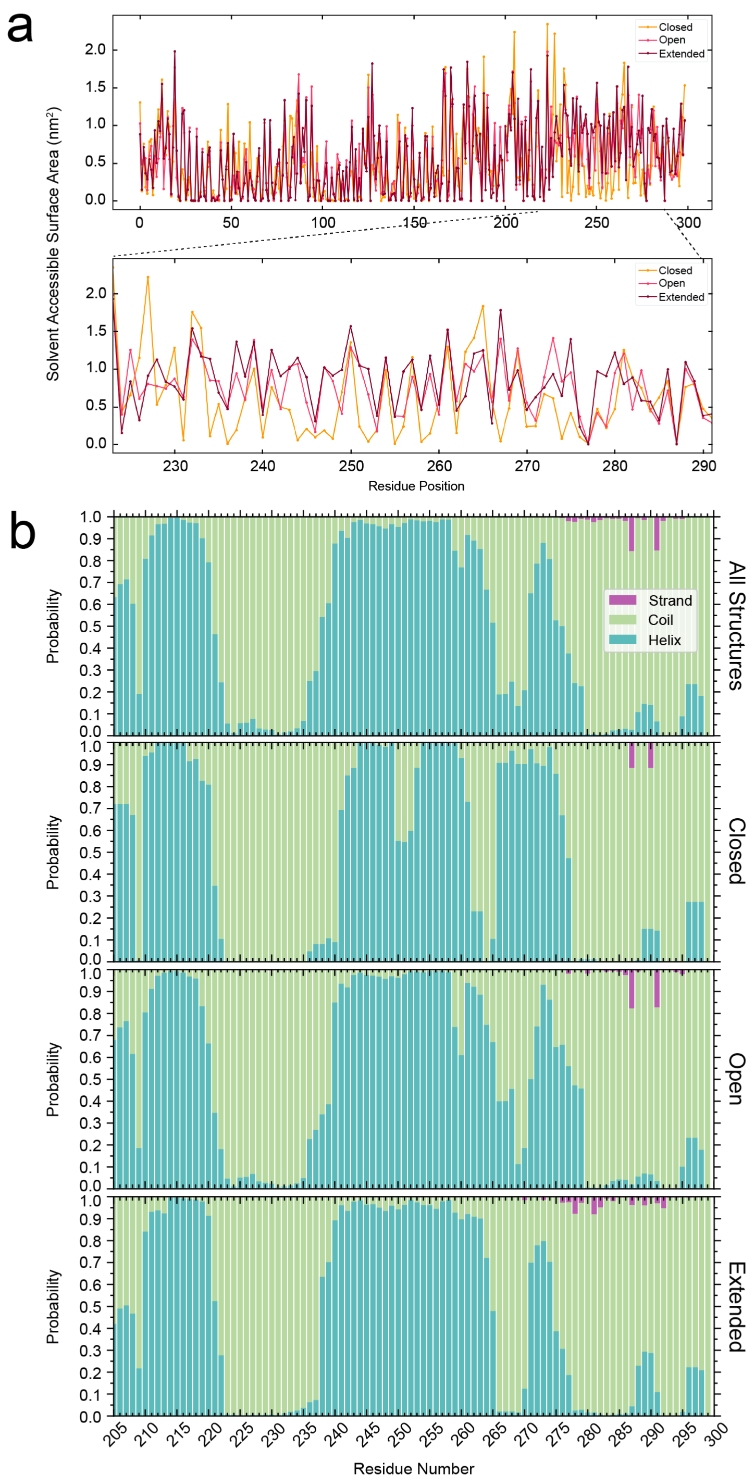


**Supplementary Figure 6. Solvent Accessible Surface Area and structural content of the C-terminal region. a.** Solvent Accessible Surface Area. Specific residues between 230 and 280 are more protected in the *extended* conformation. **b.** Secondary structure content in the C-terminal domain across the three *closed*, *open*, and *extended* subpopulations supports the presence of a structured helix with some degree of flexibility because of disordered regions.

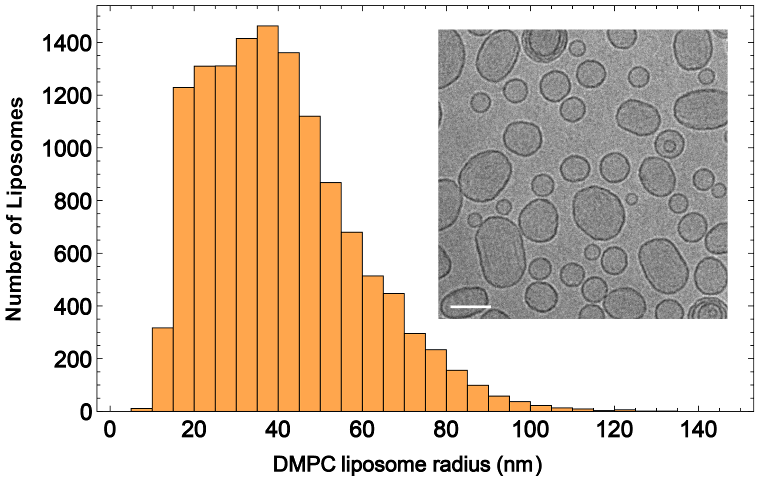


**Supplementary Figure 7**: **Size distribution of extruded liposomes determined from cryo-TEM images** (inset). The bar on the inset figure represents 100 nm.

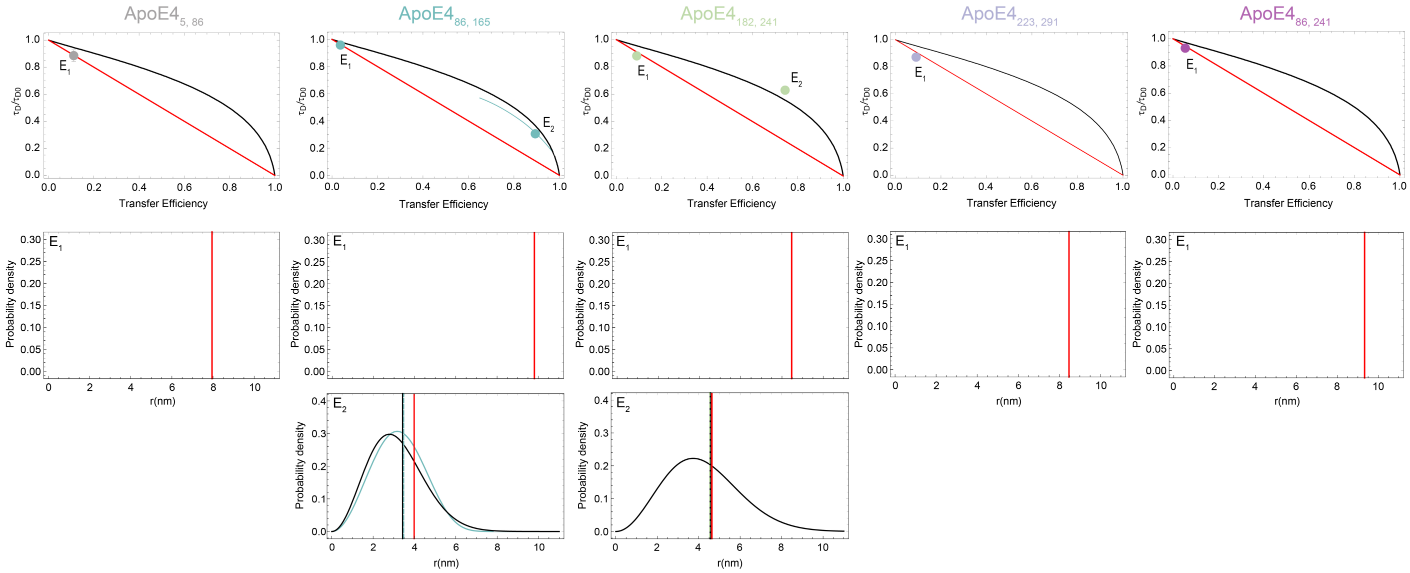


**Supplementary Figure 8**: **Lifetime *vs* transfer efficiency plot and corresponding distance distributions for each of the five ApoE4 constructs measured in the lipid-bound state** (100 µg/mL extruded liposomes, 50 mM NaPi pH 7.4). The solid red line indicates the expected result for a rigid distance. The solid black line reports the expected trend for a Gaussian distribution whose mean square end-to-end distance satisfies the measured mean transfer efficiency. The colored line represents the best fit to a wormlike chain, where both persistence length and contour length are fitted to satisfy the constraints imposed by the measured mean transfer efficiency and mean lifetime.

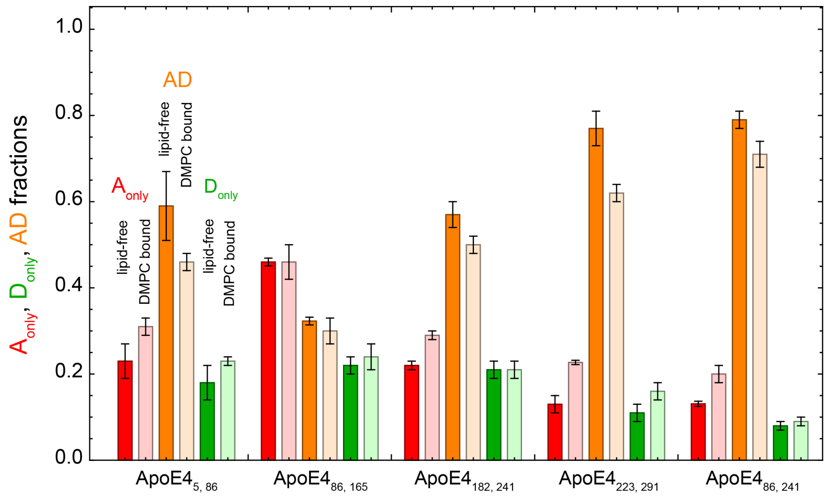


**Supplementary Figure 9**: Fractions of acceptor only (red), donor only (green), and acceptor-donor (orange) labeled molecules as identified by Pulsed Interleaved Excitation (PIE) of lipid-free (dark shaded areas) and DMPC-bound (light shaded areas) full-length ApoE constructs. Error bars are standard deviations from multiple measurements (see **Supplementary Table 14** for details). In all cases, binding to lipids results in a small decrease (up to about 15%) of the acceptor-donor labeled molecules often accompanied by a small increase in the fraction of donor and acceptor only population. These data support that only one labeled molecule of ApoE4 is binding to lipids since binding of two molecules would results in an increase of the donor-acceptor population.

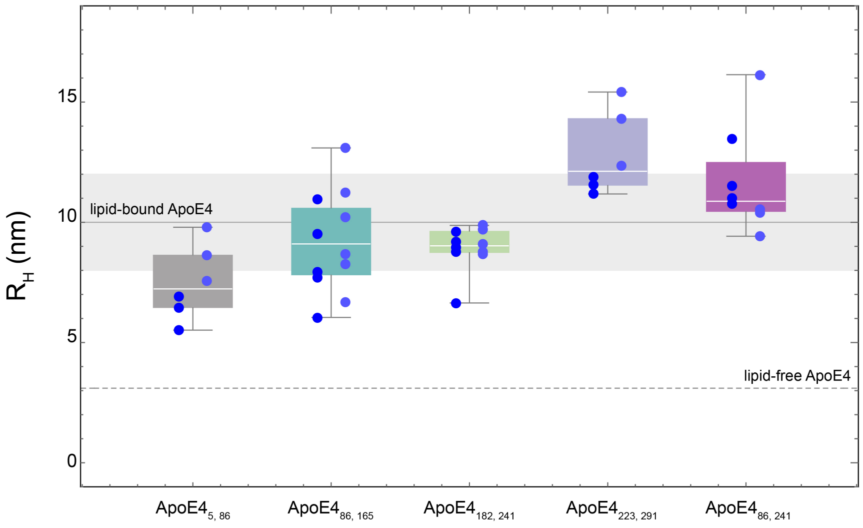


**Supplementary Figure 10**: Hydrodynamic radii ($R_H$) of lipid-bound full-length ApoE4 constructs as measured by FCS assuming one diffusing species and two triplet components. Mean $R_H$ calculated from both donor-donor and acceptor$_{PIE}$-acceptor$_{PIE}$ correlations for each full-length construct. Donor-donor and acceptor$_{PIE}$-acceptor$_{PIE}$ correlated data points are represented by dark and light blue dots, respectively. Box whisker plots report the median value as a white line, the 25% and 75% quantile, and the max and min values of the distribution for the combined donor-donor and acceptor$_{PIE}$-acceptor$_{PIE}$ datasets. Mean $R_H$ values for lipid-bound (solid gray line) and lipid-free (dashed gray line) full length ApoE4 (compare with **Supplementary Figure 11**).

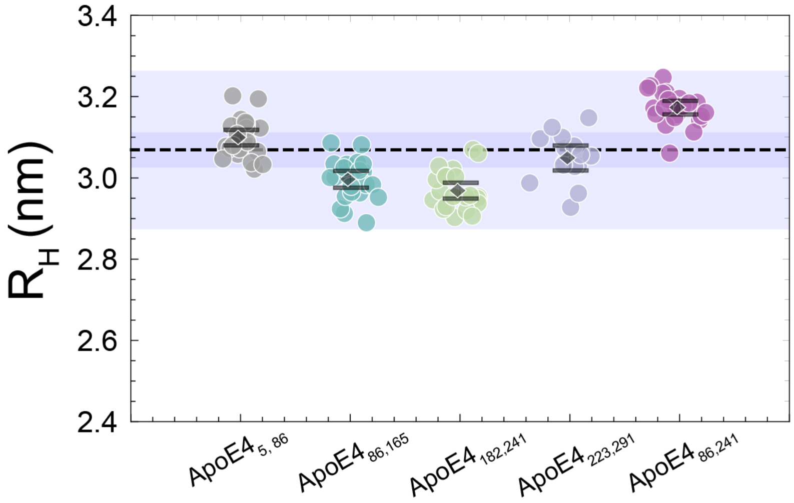


**Supplementary Figure 11**: Hydrodynamic radii of lipid-free full-length ApoE4 constructs as measured by df-FCS assuming one diffusing species and two triplet components. Variations across each construct are within a 10% of relative error and they are indistinguishable from each other according to a one-tail t-student test with significance level ≤ 0.1. Black diamonds indicate the mean $R_H$ value for each construct, and the error bars span the standard error of the mean. Dashed line denotes the overall weighted mean, and the darker shaded area spans the overall weighted SE of the mean –using individual $1/SE^2$ values as weights. Lighter shaded area denotes the overall 90% confidence interval around the overall weighted mean.

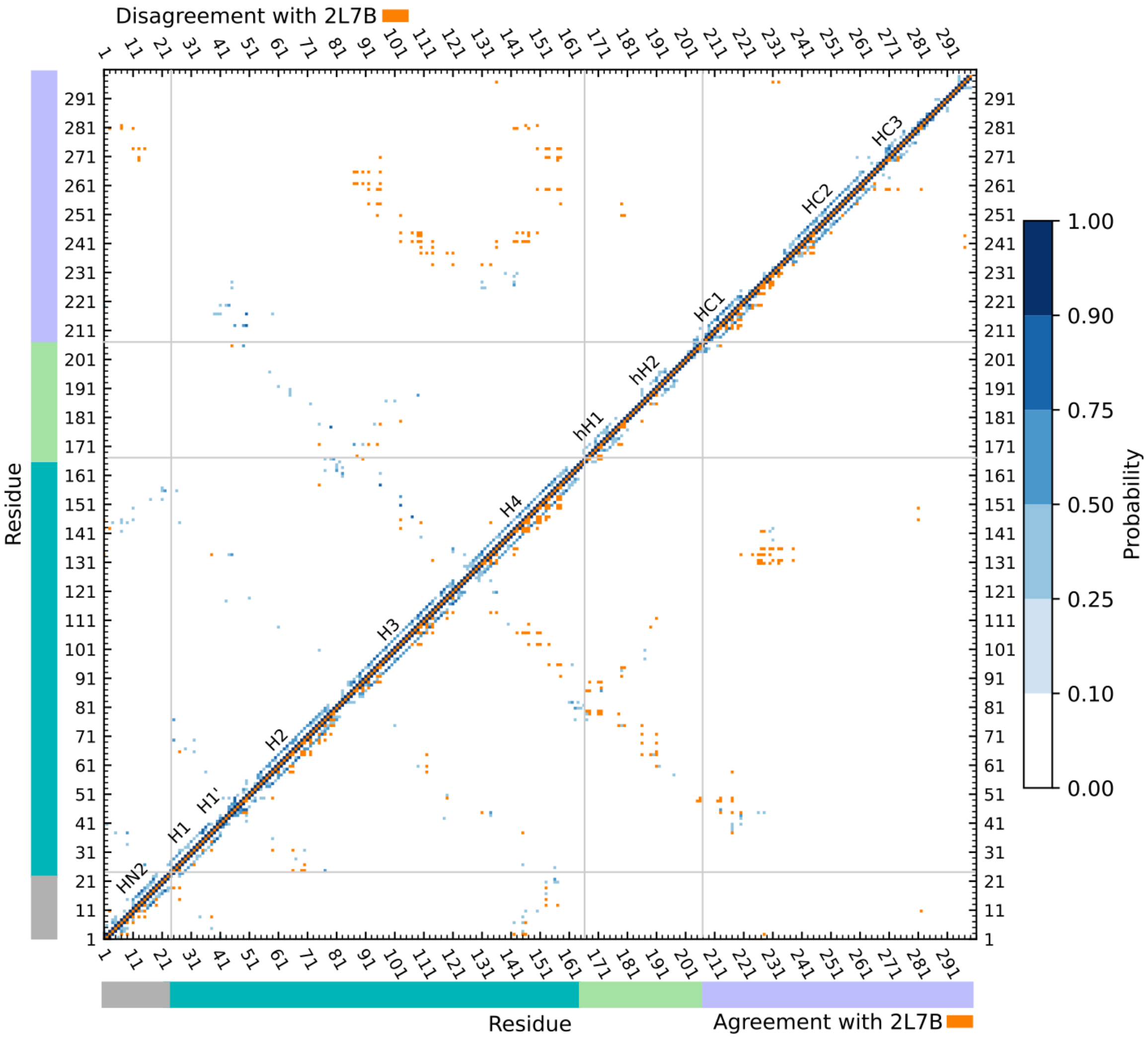


**Supplementary Figure 12.** Comparison of Folding@home simulation results ApoE3-like NMR structure, 2L7B. Displayed are the probabilities of a contact (<3 Å between any heavy atom on the two sidechains) occurring between each pairwise residue in ApoE4 according to the Folding@home equilibrium probabilities. Above the diagonal, orange marks indicate a distance predicted in the ApoE3-like NMR structure, 2L7B, but are less than 10% likely in our simulations. Below the diagonal any contacts that occur in 2L7B and are more than 10% likely in our simulations are displayed in orange.

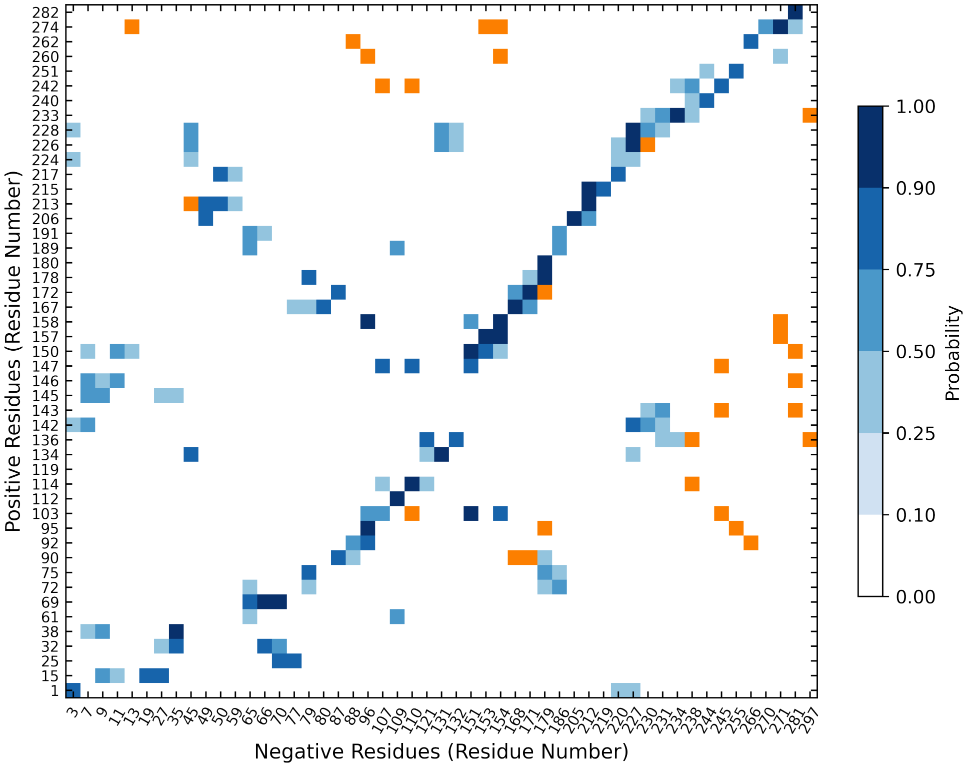


**Supplementary Figure 13.** Predicted salt bridge interactions observed in our simulations. In blue are the equilibrium probabilities of any heavy atom in the residue sidechains being within 6Å of the other sidechain. In orange are interactions predicted by the ApoE3-like NMR structure, 2L7B, which are less than 10% likely in our simulations.

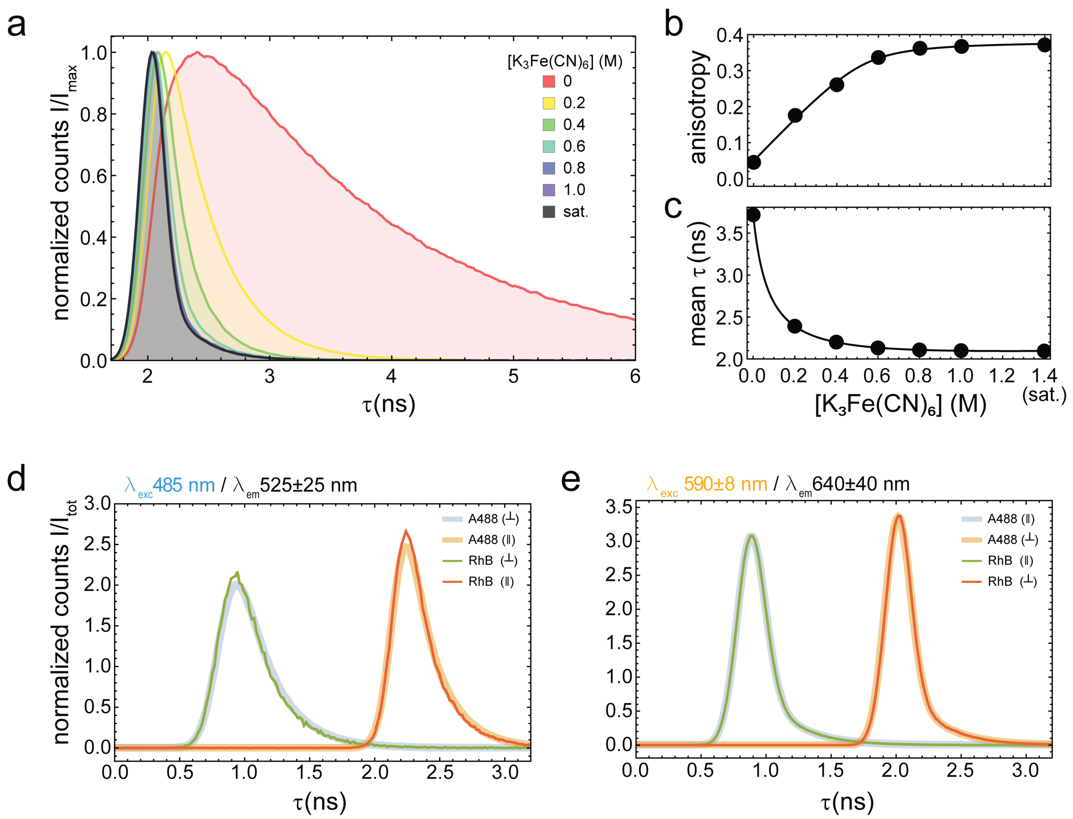


**Supplementary Figure 14**: **Instrument Response Function (IRF) determination for donor and acceptor detection. a-c.** Fluorescence lifetime histogram with normalized counts against the maximum of the distribution ($I/I_{max}$) at different concentrations of Potassium Ferrocyanide (**a**) and corresponding anisotropies (**b**) and mean photon arrival τ (**c**). **d-e**. Fluorescence lifetime histograms with normalized counts against the total area $I/I_{tot}$ for donor (**d**) and acceptor (**e**) excitations.

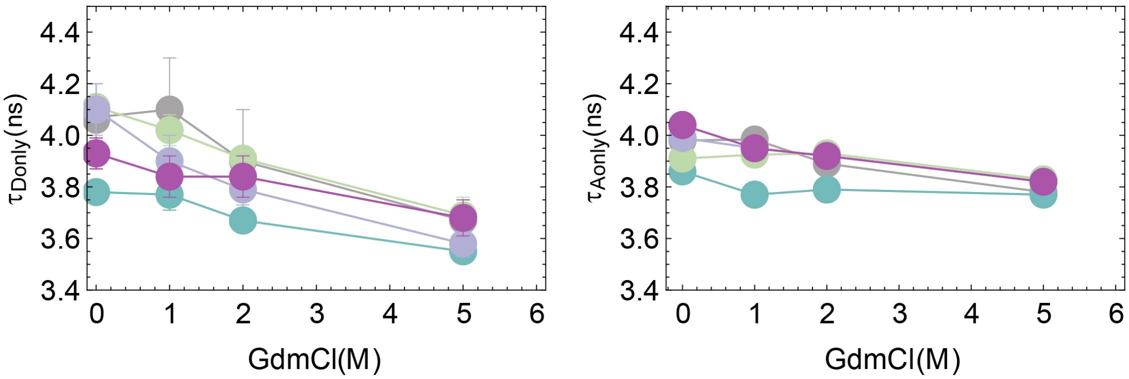


**Supplementary Figure 15**: GdmCl dependent changes in donor-only and acceptor-only lifetimes. Error bars are standard deviation from multiple measurements (**Supplementary Table 6**).

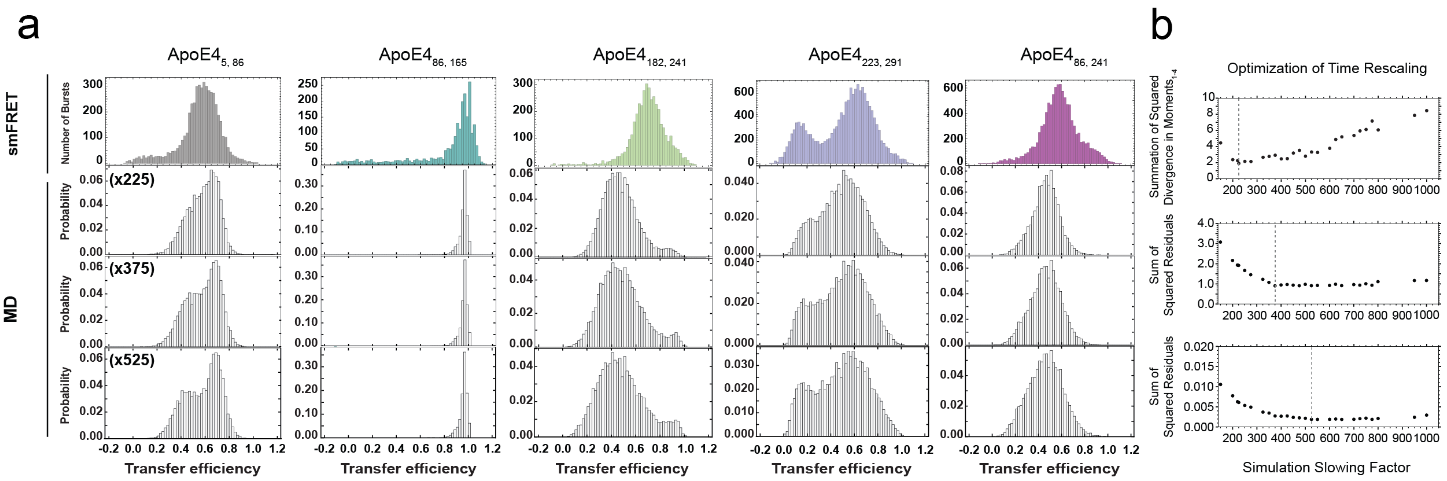


**Supplementary Figure 16. Optimization of time correction for simulations. a.** Comparison between experimental and simulated transfer efficiency histograms with different time correction factors. **b.** The time correction factors in **a.** represent the minima obtained by optimizing the factor using the sum of the residuals for the first four moments of the distributions (upper panel, time factor equals to 225), the sum of the squared residuals across each histogram normalized by maximum excursion (central panel, time factor equals to 375), and the sum of the squared residuals for only the **ApoE4$_{223,291}$** construct (lower panel, time factor equals to 525).

### Supplementary Tables

#### Supplementary Table 1. Mean transfer efficiencies, widths, and relative areas obtained from the fit of lipid-free ApoE4 smFRET histograms

**Lipid-free ApoE4$_{5, 86}$**

| GdmCl (M) | | $E_1$ | $E_2$ | $E_3$ | $W_1$ | $W_2$ | $W_3$ | $A_1$ (%) | $A_2$ (%) | $A_3$ (%) |
|---|---|---|---|---|---|---|---|---|---|---|
| 0 | n=8 | 0.21±0.05 | 0.61±0.02 | - | 0.14±0.01 | 0.118±0.008 | - | 12±5 | 88±5 | - |
| 1 | n=5 | 0.46±0.05 | 0.57±0.04 | - | 0.097±0.009 | 0.097±0.009 | - | 9±5 | 91±5 | - |
| 2* | n=5 | - | 0.5±0.1 | - | - | 0.113±0.006 | - | - | 100 | - |
| 5 | n=5 | - | 0.23±0.05 | - | - | 0.074±0.003 | - | - | 100 | - |

* fit obtained assuming a single population for the main transfer efficiency

**Lipid-free ApoE4$_{86, 165}$**

| GdmCl (M) | | $E_1$ | $E_2$ | $E_3$ | $W_1$ | $W_2$ | $W_3$ | $Asym_3$ | $A_1$ (%) | $A_2$ (%) | $A_3$ (%) |
|---|---|---|---|---|---|---|---|---|---|---|---|
| 0 | n=4 | 0.345 | 0.62 | 0.98±0.01 | 0.10 | 0.10 | 0.14±0.03 | 0.84 | 5.6±0.2 | 9±2 | 85±2 |
| 1 | n=5 | 0.345 | 0.608 | 0.978±0.005 | 0.10 | 0.10 | 0.15±0.01 | 0.84 | 6±2 | 12±3 | 82±6 |
| 2 | n=8 | 0.41±0.02 | 0.63±0.02 | 0.973±0.005 | 0.099±0.003 | 0.10 | 0.15±0.02 | 0.84 | 51±10 | 24±3 | 26±10 |
| 5 | n=6 | 0.23±0.02 | - | - | 0.076±0.009 | - | - | - | 100 | - | - |

**Lipid-free ApoE4$_{182, 241}$**

| GdmCl (M) | | $E_1$ | $E_2$ | $E_3$ | $W_1$ | $W_2$ | $W_3$ | $A_1$ (%) | $A_2$ (%) | $A_3$ (%) |
|---|---|---|---|---|---|---|---|---|---|---|
| 0 | n=6 | 0.67±0.02 | 0.83±0.02 | - | 0.078 | 0.09 | - | 61 ± 6 | 38 ± 6 | - |
| 1 | n=6 | 0.50±0.02 | 0.69±0.03 | - | 0.079 | 0.09 | - | 77 ± 6 | 23 ± 6 | - |
| 2 | n=5 | 0.44±0.02 | 0.62±0.04 | - | 0.074±0.009 | 0.09 | - | 89 ± 1 | 11 ± 1 | - |
| 5 | n=5 | 0.32±0.02 | 0.56±0.02 | - | 0.074±0.009 | 0.09 | - | 94 ± 3 | 6 ± 3 | - |

**Lipid-free ApoE4$_{223, 291}$**

| GdmCl (M) | | $E_1$ | $E_2$ | $E_3$ | $W_1$ | $W_2$ | $W_3$ | $A_1$ (%) | $A_2$ (%) | $A_3$ (%) |
|---|---|---|---|---|---|---|---|---|---|---|
| 0 | n=12 | 0.13±0.04 | 0.61±0.02 | 0.851±0.003 | 0.105±0.004 | 0.13 | 0.104±0.001 | 27 ± 4 | 58 ± 4 | 15 ± 5 |
| 1 | n=6 | 0.15±0.06 | 0.46±0.02 | 0.85 | 0.104 | 0.095±0.003 | 0.104 | 10 ± 6 | 85 ± 6 | 5 ± 1 |
| 2 | n=6 | - | 0.37±0.02 | - | - | 0.093±0.009 | - | - | 100 | - |
| 5 | n=5 | - | 0.27±0.02 | - | - | 0.079±0.006 | - | - | 100 | - |

**Lipid-free ApoE4$_{86, 241}$**

| GdmCl (M) | | $E_1$ | $E_2$ | $E_3$ | $W_1$ | $W_2$ | $W_3$ | $A_1$ (%) | $A_2$ (%) | $A_3$ (%) |
|---|---|---|---|---|---|---|---|---|---|---|
| 0 | n=5 | 0.24±0.01 | 0.59±0.02 | 0.87±0.02 | 0.12 | 0.113±0.007 | 0.09±0.01 | 7 ± 1 | 81 ± 1 | 12 ± 1 |
| 1 | n=5 | 0.16±0.02 | 0.40±0.03 | 0.80 | 0.06±0.02 | 0.089±0.003 | 0.12 | 2.2 ± 0.3 | 92 ± 1 | 6 ± 1 |
| 2 | n=4 | - | 0.25±0.04 | - | - | 0.106±0.008 | - | - | 100 | - |
| 5 | n=4 | - | 0.10±0.03 | - | - | 0.06±0.01 | - | - | 100 | - |

**Supplementary Table 2. Folding free energy and midpoint of the transition determined from folding equilibrium analysis**

| 3-state model | | | | | |
|---|---|---|---|---|---|
| | $ApoE4_{5,86}$ | $ApoE4_{86,165}$ | $ApoE4_{182,241}$ | $ApoE4_{223,291}$ | $ApoE4_{86,241}$ |
| $c_{1/2\ UN1}$ (M) | 2.06±0.01 | 1.91±0.015 | 1.9±0.2 | -0.8±0.1 | -1.00±0.07 |
| $\Delta G^0_{UN1}$ (RT) | -5.2±0.2 | -8.3±0.42 | -8±7 | 1.15±0.08 | 1.47±0.04 |
| $c_{1/2\ N1N2}$ (M) | -0.6±0.4 | 3±0.2 | -0.3±0.1 | 1.40±0.09 | -1.2±0.7 |
| $\Delta G^0_{N1N2}$ (RT) | 2.1±0.4 | 2.7±0.2 | 1.0±0.2 | 1.6±0.1 | 0.7±0.1 |

| 2-state model | |
|---|---|
| | $ApoE4_{5,86}$ |
| $c_{1/2\ UN1}$ (M) | 2.07±0.03 |
| $\Delta G^0_{UN1}$ (RT) | -5.4±0.4 |
| $c_{1/2\ N1N2}$ (M) | -0.41±0.06 |
| $\Delta G^0_{N1N2}$ (RT) | 1.76±0.08 |

**Supplementary Table 3. Folding of ApoE4 as reported in literature.**

| Region | Residues | $c_{1/2}$ (M) | $\Delta G^0_{UN}$ (kcal $mol^{-1}$) | reference |
|---|---|---|---|---|
| Four-helix bundle | 1-183 | 2 | - | Weers et al. 2003[27] |
| Four-helix bundle | 1-191 | 2 | - | Morrow et al. 2000[25] |
| N-terminal domain | full-length | 2.6 ± 1.3 | 5.25 ± 0.20 | Dolai et al. [26] |
| C-terminal domain | full-length | 0.73 ± 0.07 | 3.29 ± 0.20 | Dolai et al. [26] |
| N-terminal domain (1-191) | fragment | - | 4.56 ± 0.03 | Dolai et al. [26] |
| C-terminal domain (192-299) | fragment | - | 1.58 ± 0.08 | Dolai et al. [26] |

**Supplementary Table 4. Mean transfer efficiencies, widths, and relative areas obtained from the fit of lipid-free ApoE4 smFRET histograms**

| Lipid-bound ApoE4 | | | | | | | | | | |
|---|---|---|---|---|---|---|---|---|---|---|
| Construct | | $E_1$ | $E_2$ | $E_3$ | $W_1$ | $W_2$ | $W_3$ | $A_1$ (%) | $A_2$ (%) | $A_3$ (%) |
| 5, 86 | n=3 | 0.111±0.006 | - | - | 0.073±0.002 | - | - | 100 | - | - |
| 86, 165 | n=6 | 0.037±0.006 | 0.894±0.004 | - | 0.17±0.01 | 0.080±0.009 | - | 51 ± 16 | 49 ± 16 | - |
| 182, 241 | n=4 | 0.09±0.01 (1.72±0.07) | 0.744 | - | 0.21±0.03 | 0.137 | - | 91 ± 2 | 9 ± 2 | - |
| 223, 291 | n=3 | 0.091±0.008 | - | - | 0.072±0.002 | - | - | 100 | - | - |
| 86, 241 | n=3 | 0.054±0.003 (1.57±0.04) | - | - | 0.215±0.007 | - | - | 100 | - | - |

()=Asymmetry of lognormal distribution

**Supplementary Table 5. Correction parameters for anisotropy determinations.**

| Excitation | Detection | $L_s$ | $L_p$ | $G$ |
|---|---|---|---|---|
| Donor | Acceptor | 0.036 | 0.218 | 0.776 |
| | Donor | 0.072 | 0.126 | 0.992 |
| Acceptor | Acceptor | 0.042 | 0.169 | 1.29 |

**Supplementary Table 6. Subpopulation-specific analysis of time-resolved fluorescence decays for lipid-free ApoE4**

**$ApoE4_{5, 86}$**

| GdmCl (M) | | | $\tau_{Donly}$ (ns) | $\tau_{rot\text{-}Donly}$(ns) | $\tau_{DA}$(ns) | $\tau_{rot\text{-}DA}$(ns) | $\tau_{FRET}$(ns) | $\tau_{Aonly}$(ns) | $\tau_{rot\text{-}Aonly}$(ns) |
|---|---|---|---|---|---|---|---|---|---|
| 0 | $E_1$ | n=6 | 4.06±0.08 | 0.43±0.07 | 3.28±0.08 | 0.7±0.1 | 17±3 | 3.98±0.02 | 0.54±0.04 |
| 0 | $E_2$ | n=6 | 4.06±0.08 | 0.44±0.06 | 2.13±0.07 | 0.62±0.06 | 4.5±0.3 | 3.98±0.02 | 0.53±0.04 |
| 1 | $E_1$ | n=3 | 4.1±0.2 | 0.52±0.06 | 2.28±0.04 | 0.58±0.02 | 5.1±0.4 | 3.984±0.004 | 0.7±0.1 |
| 2 | $E_1$ | n=3 | 3.9±0.2 | 0.51±0.01 | 2.45±0.03 | 0.61±0.02 | 6.6±0.6 | 3.89±0.03 | 0.79±0.03 |
| 5 | $E_1$ | n=3 | 3.67±0.08 | 0.60±0.03 | 2.90±0.01 | 0.58±0.01 | 16±2 | 3.78±0.01 | 0.99±0.09 |

**$ApoE4_{86, 165}$**

| GdmCl (M) | | | $\tau_{Donly}$ (ns) | $\tau_{rot\text{-}Donly}$(ns) | $\tau_{DA}$(ns) | $\tau_{rot\text{-}DA}$(ns) | $\tau_{FRET}$(ns) | $\tau_{Aonly}$(ns) | $\tau_{rot\text{-}Aonly}$(ns) |
|---|---|---|---|---|---|---|---|---|---|
| 0 | $E_3$ | n=3 | 3.78±0.03 | 0.32±0.04 | 0.68±0.05 | 0.225±0.008 | 0.83±0.07 | 3.86±0.01 | 0.40±0.01 |
| 1 | $E_3$ | n=3 | 3.77±0.06 | 0.48±0.02 | 0.82±0.02 | 0.30±0.01 | 1.05±0.03 | 3.77±0.02 | 0.51±0.02 |
| 1 | $E_2$ | n=3 | 3.77±0.06 | 0.48±0.02 | 2.68±0.01 | 0.62±0.09 | 9.3±0.4 | 3.77±0.02 | 0.51±0.02 |
| 2 | $E_3$ | n=2 | 3.67±0.02 | 0.61±0.03 | 0.86±0.09 | 0.29±0.01 | 1.1±0.2 | 3.79±0.02 | 0.75±0.08 |
| 2 | $E_2$ | n=2 | 3.67±0.02 | 0.61±0.03 | 2.30±0.04 | 0.578±0.009 | 6.2±0.3 | 3.79±0.02 | 0.75±0.08 |
| 2 | $E_1$ | n=2 | 3.67±0.02 | 0.61±0.03 | 2.782±0.007 | 0.64±0.02 | 11.5±0.2 | 3.79±0.02 | 0.75±0.08 |
| 5 | $E_1$ | n=3 | 3.55±0.03 | 0.724±0.009 | 3.030±0.006 | 0.56±0.01 | 21±1 | 3.77±0.01 | 1.03±0.05 |

**$ApoE4_{182, 241}$**

| GdmCl (M) | | | $\tau_{Donly}$ (ns) | $\tau_{rot\text{-}Donly}$(ns) | $\tau_{DA}$(ns) | $\tau_{rot\text{-}DA}$(ns) | $\tau_{FRET}$(ns) | $\tau_{Aonly}$(ns) | $\tau_{rot\text{-}Aonly}$(ns) |
|---|---|---|---|---|---|---|---|---|---|
| 0 | $E_2$ | n=4 | 4.11±0.04 | 0.62±0.02 | 1.24±0.07 | 0.31±0.01 | 1.8±0.1 | 3.91±0.02 | 0.52±0.03 |
| 0 | $E_1$ | n=4 | 4.11±0.04 | 0.62±0.02 | 2.07±0.04 | 0.52±0.02 | 6.2±0.2 | 3.91±0.02 | 0.52±0.03 |
| 1 | $E_2$ | n=3 | 4.02±0.05 | 0.68±0.06 | 2.47±0.02 | 0.65±0.05 | 6.4±0.2 | 3.925±0.008 | 0.9±0.1 |
| 1 | $E_1$ | n=3 | 4.02±0.05 | 0.68±0.06 | 2.63±0.01 | 0.74±0.04 | 7.6±0.2 | 3.925±0.008 | 0.9±0.1 |
| 2 | $E_2$ | n=3 | 3.91±0.04 | 0.71±0.05 | 2.55±0.03 | 0.63±0.03 | 7.3±0.3 | 3.93±0.03 | 1.0±0.1 |
| 2 | $E_1$ | n=3 | 3.91±0.04 | 0.71±0.05 | 2.77±0.02 | 0.775±0.009 | 9.5±0.3 | 3.93±0.03 | 1.0±0.1 |
| 5 | $E_1$ | n=3 | 3.69±0.07 | 0.75±0.04 | 2.85±0.01 | 0.72±0.01 | 13±1 | 3.83±0.04 | 1.15±0.05 |

**$ApoE4_{223, 291}$**

| GdmCl (M) | | | $\tau_{Donly}$ (ns) | $\tau_{rot\text{-}Donly}$(ns) | $\tau_{DA}$(ns) | $\tau_{rot\text{-}DA}$(ns) | $\tau_{FRET}$(ns) | $\tau_{Aonly}$(ns) | $\tau_{rot\text{-}Aonly}$(ns) |
|---|---|---|---|---|---|---|---|---|---|
| 0 | $E_3$ | n=4 | 4.1±0.1 | 0.6±0.2 | 1.59±0.08 | 0.40±0.04 | 2.6±0.2 | 3.99±0.02 | 0.68±0.02 |
| 0 | $E_2$ | n=4 | 4.1±0.1 | 0.6±0.2 | 2.33±0.01 | 0.59±0.03 | 5.4±0.2 | 3.99±0.02 | 0.68±0.02 |
| 0 | $E_1$ | n=4 | 4.1±0.1 | 0.6±0.2 | 3.35±0.02 | 0.6±0.1 | 18±2 | 3.99±0.02 | 0.68±0.02 |
| 1 | $E_2$ | n=3 | 3.9±0.1 | 0.7±0.1 | 2.65±0.03 | 0.74±0.02 | 8.3±0.5 | 3.95±0.02 | 0.92±0.03 |
| 2 | $E_2$ | n=3 | 3.79±0.06 | 0.60±0.05 | 2.793±0.006 | 0.744±0.007 | 10.6±0.5 | 3.92±0.01 | 0.99±0.02 |
| 5 | $E_2$ | n=3 | 3.58±0.05 | 0.69±0.01 | 2.92±0.02 | 0.71±0.02 | 16±1 | 3.82±0.03 | 1.2±0.1 |

**$ApoE4_{86, 241}$**

| GdmCl (M) | | | $\tau_{Donly}$ (ns) | $\tau_{rot\text{-}Donly}$(ns) | $\tau_{DA}$(ns) | $\tau_{rot\text{-}DA}$(ns) | $\tau_{FRET}$(ns) | $\tau_{Aonly}$(ns) | $\tau_{rot\text{-}Aonly}$(ns) |
|---|---|---|---|---|---|---|---|---|---|
| 0 | $E_3$ | n=3 | 3.93±0.06 | 0.51±0.08 | 1.33±0.04 | 0.37±0.03 | 2.01±0.09 | 4.04±0.01 | 0.34±0.01 |
| 0 | $E_2$ | n=3 | 3.93±0.06 | 0.51±0.08 | 2.45±0.01 | 0.64±0.03 | 6.5±0.2 | 4.04±0.01 | 0.34±0.01 |
| 0 | $E_1$ | n=3 | 3.93±0.06 | 0.51±0.08 | 3.30±0.04 | 0.70±0.02 | 21±2 | 4.04±0.01 | 0.34±0.01 |
| 1 | $E_2$ | n=3 | 3.84±0.08 | 0.53±0.04 | 2.92±0.02 | 0.63±0.02 | 12.2±0.6 | 3.95±0.02 | 0.8±0.2 |
| 2 | $E_2$ | n=3 | 3.84±0.08 | 0.56±0.02 | 3.18±0.01 | 0.65±0.02 | 18.5±1.9 | 3.92±0.01 | 0.89±0.06 |
| 5 | $E_2$ | n=3 | 3.68±0.07 | 0.537±0.005 | 3.32±0.02 | 0.64±0.01 | 34±6 | 3.82±0.03 | 0.95±0.03 |

**Supplementary Table 7. Subpopulation-specific analysis of time-resolved fluorescence decays for lipid-bound ApoE4**

| Construct | E | | $\tau_{Donly}$ (ns) | $\tau_{rot\text{-}Donly}$(ns) | $\tau_{DA}$(ns) | $\tau_{rot\text{-}DA}$(ns) | $\tau_{FRET}$(ns) | $\tau_{Aonly}$(ns) | $\tau_{rot\text{-}Aonly}$(ns) |
|---|---|---|---|---|---|---|---|---|---|
| $ApoE4_{5,\ 86}$ | $E_1$ | n=3 | 4.2±0.2 | 0.31±0.02 | 3.72±0.02 | 0.43±0.01 | 33±12 | 4.01±0.07 | 0.46±0.07 |
| $ApoE4_{86,\ 165}$ | $E_1$ | n=5 | 4.06±0.05 | 0.33±0.02 | 3.89±0.04 | 0.33±0.03 | 93±35 | 4.04±0.06 | 0.29±0.06 |
| $ApoE4_{86,\ 165}$ | $E_2$ | n=5 | 4.06±0.05 | 0.33±0.02 | 1.24±0.06 | 0.31±0.02 | 1.79±0.12 | 4.04±0.06 | 0.29±0.06 |
| $ApoE4_{182,\ 241}$ | $E_1$ | n=4 | 4.13±0.06 | 0.29±0.02 | 3.64±0.04 | 0.34±0.03 | 31±4 | 4.07±0.06 | 0.33±0.04 |
| $ApoE4_{182,\ 241}$ | $E_2$ | n=4 | 4.13±0.06 | 0.29±0.02 | 2.6±0.1 | 0.40±0.02 | 7.0±0.7 | 4.07±0.06 | 0.33±0.04 |
| $ApoE4_{223,\ 291}$ | $E_1$ | n=3 | 4.21±0.09 | 0.30±0.01 | 3.67±0.02 | 0.37±0.03 | 29±4 | 4.27±0.03 | 0.43±0.03 |
| $ApoE4_{86,\ 241}$ | $E_1$ | n=4 | 4.08±0.04 | 0.31±0.04 | 3.8±0.1 | 0.37±0.04 | 52±25 | 4.10±0.04 | 0.16±0.04 |

**Supplementary Table 8. Steady-state and time resolved anisotropies in aqueous buffer conditions.**

| Construct | E | | $r_{ss\text{-}Donly}$ | $r_{\infty\text{-}Donly}$ | $\tau_{rot\text{-}Donly}$ (ns) | $r_{ss\text{-}Aonly}$ | $r_{\infty\text{-}Aonly}$ | $\tau_{rot\text{-}Aonly}$ (ns) | $r_{ss\text{-}A(D)}$ | $r_{\infty\text{-}A(D)}$ |
|---|---|---|---|---|---|---|---|---|---|---|
| $ApoE4_{5,\ 86}$ | $E_2$ | n=6 | 0.09±0.01 | 0.09±0.02 | 0.47±0.04 | 0.08±0.01 | 0.07±0.03 | 0.45±0.06 | 0.062±0.004 | 0.06±0.01 |
| $ApoE4_{5,\ 86}$ | $E_1$ | n=6 | 0.09±0.01 | 0.09±0.02 | 0.47±0.04 | 0.08±0.01 | 0.07±0.03 | 0.45±0.06 | 0.087±0.007 | 0.10±0.02 |
| $ApoE4_{86,\ 165}$ | $E_3$ | n=5 | 0.136±0.003 | 0.143±0.009 | 0.36±0.06 | 0.097±0.005 | 0.102±0.003 | 0.37±0.01 | 0.049±0.003 | 0.043±0.006 |
| $ApoE4_{182,\ 241}$ | $E_2$ | n=4 | 0.131±0.005 | 0.15±0.01 | 0.50±0.06 | 0.099±0.009 | 0.10±0.01 | 0.43±0.04 | 0.060±0.006 | 0.061±0.005 |
| $ApoE4_{182,\ 241}$ | $E_1$ | n=4 | 0.131±0.005 | 0.15±0.01 | 0.50±0.06 | 0.099±0.009 | 0.10±0.01 | 0.43±0.04 | 0.057±0.003 | 0.064±0.004 |
| $ApoE4_{223,\ 291}$ | $E_3$ | n=4 | 0.13±0.02 | 0.16±0.02 | 0.48±0.06 | 0.13±0.01 | 0.146±0.009 | 0.50±0.05 | 0.065±0.004 | 0.069±0.009 |
| $ApoE4_{223,\ 291}$ | $E_2$ | n=4 | 0.13±0.02 | 0.16±0.02 | 0.48±0.06 | 0.13±0.01 | 0.146±0.009 | 0.50±0.05 | 0.063±0.001 | 0.060±0.004 |
| $ApoE4_{223,\ 291}$ | $E_1$ | n=4 | 0.13±0.02 | 0.16±0.02 | 0.48±0.06 | 0.13±0.01 | 0.146±0.009 | 0.50±0.05 | 0.112±0.007 | 0.15±0.02 |
| $ApoE4_{86,\ 241}$ | $E_3$ | n=4 | 0.142±0.007 | 0.146±0.003 | 0.54±0.01 | 0.142±0.003 | 0.148±0.007 | 0.42±0.05 | 0.063±0.003 | 0.058±0.006 |
| $ApoE4_{86,\ 241}$ | $E_2$ | n=4 | 0.142±0.007 | 0.146±0.003 | 0.54±0.01 | 0.142±0.003 | 0.148±0.007 | 0.42±0.05 | 0.0604±0.0007 | 0.069±0.003 |
| $ApoE4_{86,\ 241}$ | $E_1$ | n=4 | 0.142±0.007 | 0.146±0.003 | 0.54±0.01 | 0.142±0.003 | 0.148±0.007 | 0.42±0.05 | 0.080±0.004 | 0.08±0.01 |

**Supplementary Table 10. Steady-state and time resolved anisotropies for lipid-bound ApoE4**

| Construct | E | | $r_{ss\text{-}Donly}$ | $r_{\infty\text{-}Donly}$ | $\tau_{rot\text{-}Donly}$ (ns) | $r_{ss\text{-}Aonly}$ | $r_{\infty\text{-}Aonly}$ | $\tau_{rot\text{-}Aonly}$ (ns) | $r_{ss\text{-}A(D)}$ | $r_{\infty\text{-}A(D)}$ |
|---|---|---|---|---|---|---|---|---|---|---|
| $ApoE4_{5,\ 86}$ | $E_1$ | n=3 | 0.078±0.002 | 0.10±0.02 | 0.34±0.02 | 0.07±0.01 | 0.053±0.007 | 0.66±0.02 | 0.121±0.002 | 0.14±0.01 |
| $ApoE4_{86,\ 165}$ | $E_1$ | n=5 | 0.16±0.01 | 0.149±0.009 | 0.64±0.09 | 0.10±0.02 | 0.11±0.03 | 0.57±0.02 | 0.17±0.01 | 0.24±0.03 |
| $ApoE4_{86,\ 165}$ | $E_2$ | n=5 | 0.16±0.01 | 0.149±0.009 | 0.64±0.09 | 0.10±0.02 | 0.11±0.03 | 0.57±0.02 | 0.050±0.006 | 0.051±0.004 |
| $ApoE4_{182,\ 241}$ | $E_1$ | n=4 | 0.134±0.009 | 0.14±0.01 | 0.47±0.08 | 0.139±0.006 | 0.147±0.009 | 0.71±0.04 | 0.131±0.003 | 0.17±0.03 |
| $ApoE4_{182,\ 241}$ | $E_2$ | n=4 | 0.134±0.009 | 0.14±0.01 | 0.47±0.08 | 0.139±0.006 | 0.147±0.009 | 0.71±0.04 | 0.083±0.006 | 0.085±0.004 |
| $ApoE4_{223,\ 291}$ | $E_1$ | n=3 | 0.13±0.01 | 0.15±0.01 | 0.44±0.04 | 0.14±0.01 | 0.16±0.01 | 0.75±0.04 | 0.141±0.002 | 0.208±0.009 |
| $ApoE4_{86,\ 241}$ | $E_1$ | n=4 | 0.14±0.02 | 0.14±0.02 | 0.54±0.06 | 0.17±0.02 | 0.20±0.01 | 0.61±0.04 | 0.152±0.003 | 0.22±0.01 |

**Supplementary Table 11. Estimates of deviations in the orientational $\kappa^2$ factor in aqueous buffer conditions.**

| Construct | E | $\kappa^2_{min}$ | $\kappa^2_{max}$ | $\kappa^2_{mean}$ | precision | accuracy | Corrected $R_0$ |
|---|---|---|---|---|---|---|---|
| $ApoE4_{5,\ 86}$ | $E_2$ | 0.36 (0.35) | 1.75 (1.77) | 0.79 (0.79) | 0.01 (0.01) | 0.06 (0.07) | 5.55 |
| $ApoE4_{5,\ 86}$ | $E_1$ | 0.36 (0.35) | 1.75 (1.77) | 0.88 (0.91) | 0.03 (0.03) | 0.07 (0.07) | 5.66 |
| $ApoE4_{86,\ 165}$ | $E_3$ | 0.29 (0.30) | 2.17 (2.12) | 0.73 (0.74) | 0.006 (0.002) | 0.07 (0.07) | 5.48 |
| $ApoE4_{182,\ 241}$ | $E_2$ | 0.29 (0.30) | 2.22 (2.09) | 0.76 (0.76) | 0.0002 (0.002) | 0.07 (0.07) | 5.52 |
| $ApoE4_{182,\ 241}$ | $E_1$ | 0.29 (0.30) | 2.22 (2.09) | 0.77 (0.76) | 0.0008 (0.0008) | 0.07 (0.07) | 5.53 |
| $ApoE4_{223,\ 291}$ | $E_3$ | 0.24 (0.28) | 2.35 (2.13) | 0.76 (0.76) | 0.005 (0.0006) | 0.08 (0.08) | 5.52 |
| $ApoE4_{223,\ 291}$ | $E_2$ | 0.24 (0.28) | 2.35 (2.13) | 0.75 (0.76) | 0.008 (0.001) | 0.08 (0.08) | 5.51 |
| $ApoE4_{223,\ 291}$ | $E_1$ | 0.24 (0.28) | 2.35 (2.13) | 0.98 (0.88) | 0.03 (0.02) | 0.1 (0.08) | 5.76 |
| $ApoE4_{86,\ 241}$ | $E_3$ | 0.25 (0.26) | 2.26 (2.23) | 0.74 (0.76) | 0.007 (0.004) | 0.08 (0.08) | 5.49 |
| $ApoE4_{86,\ 241}$ | $E_2$ | 0.25 (0.26) | 2.26 (2.23) | 0.76 (0.75) | 0.004 (0.005) | 0.08 (0.08) | 5.52 |
| $ApoE4_{86,\ 241}$ | $E_1$ | 0.25 (0.26) | 2.26 (2.23) | 0.78 (0.79) | 0.0004 (0.0009) | 0.08 (0.08) | 5.54 |

**Supplementary Table 12. Estimates of deviations in the orientational $\kappa^2$ factor for lipid-bound ApoE4**

| Construct | E | $\kappa^2_{min}$ | $\kappa^2_{max}$ | $\kappa^2_{mean}$ | precision | accuracy | Corrected $R_0$ |
|---|---|---|---|---|---|---|---|
| $ApoE4_{5, 86}$ | $E_1$ | 0.37 (0.37) | 1.78 (1.66) | 0.85 (0.86) | 0.03 (0.03) | 0.05 (0.06) | 5.62 |
| $ApoE4_{86, 165}$ | $E_1$ | 0.28 (0.28) | 2.23 (2.28) | 0.92 (0.95) | 0.03 (0.03) | 0.07 (0.08) | 5.70 |
| $ApoE4_{86, 165}$ | $E_2$ | 0.28 (0.28) | 2.23 (2.28) | 0.74 (0.74) | 0.005 (0.005) | 0.07 (0.07) | 5.49 |
| $ApoE4_{182, 241}$ | $E_1$ | 0.26 (0.26) | 2.16 (2.21) | 0.98 (0.92) | 0.03 (0.02) | 0.09 (0.09) | 5.76 |
| $ApoE4_{182, 241}$ | $E_2$ | 0.26 (0.26) | 2.22 (2.21) | 0.80 (0.79) | 0.002 (0.003) | 0.08 (0.08) | 5.56 |
| $ApoE4_{223, 291}$ | $E_1$ | 0.24 (0.27) | 2.30 (2.15) | 0.99 (0.99) | 0.03 (0.03) | 0.09 (0.09) | 5.77 |
| $ApoE4_{86, 241}$ | $E_1$ | 0.22 (0.24) | 2.29 (2.25) | 1.01 (0.99) | 0.03 (0.03) | 0.10 (0.10) | 5.79 |

()=values estimated from steady-state anisotropies.

**Supplementary Table 13. $\sigma^2$ values (Eq. 18) for ApoE4 under aqueous buffer and lipid-bound conditions**

| Construct | | | $E_{1\text{-Measured}}$ | $E_{1\text{-Gaussian}}$ | $E_{2\text{-Measured}}$ | $E_{2\text{-Gaussian}}$ | $E_{3\text{-Measured}}$ | $E_{3\text{-Gaussian}}$ |
|---|---|---|---|---|---|---|---|---|
| $ApoE4_{5, 86}$ | lipid-free | n=6 | 0.02±0.01 | 0.08±0.02 | 0.055±0.007 | 0.107±0.004 | - | - |
| | DMPC | n=3 | 0.01±0.01 | 0.050±0.002 | - | - | - | - |
| $ApoE4_{86, 165}$ | lipid-free | n=3 | 0.14±0.03 | 0.115 | 0.09±0.01 | 0.107 | 0.0010±0.0006 | 0.0003±0.0004 |
| | DMPC | n=6 | 0.003±0.003 | 0.017±0.003 | 0.021±0.001 | 0.028±0.001 | - | - |
| $ApoE4_{182, 241}$ | lipid-free | n=4 | 0.061±0.002 | 0.089±0.002 | 0.022±0.003 | 0.035±0.007 | - | - |
| | DMPC | n=4 | 0 | 0.039±0.005 | 0.095±0.005 | 0.0769 | - | - |
| $ApoE4_{223, 291}$ | lipid-free | n=4 | 0.01±0.01 | 0.0761±0.0003 | 0.075±0.007 | 0.104±0.001 | 0.035±0.004 | 0.040±0.004 |
| | DMPC | n=3 | 0 | 0.041±0.003 | - | - | - | - |
| $ApoE4_{86, 241}$ | lipid-free | n=3 | 0.07±0.01 | 0.097 | 0.091±0.003 | 0.1094±0.0003 | 0.0255±0.0009 | 0.0297±0.0007 |
| | DMPC | n=3 | 0 | 0.025±0.001 | - | - | - | - |

**Supplementary Table 14. Stoichiometry ratios for ApoE4 under aqueous buffer and lipid-bound conditions.**

| Construct | | | $A_{only}$ | DA | $D_{only}$ |
|---|---|---|---|---|---|
| $ApoE4_{5, 86}$ | lipid-free | n=6 | 0.23±0.04 | 0.59±0.08 | 0.18±0.04 |
| | DMPC | n=3 | 0.31±0.02 | 0.46±0.02 | 0.23±0.01 |
| $ApoE4_{86, 165}$ | lipid-free | n=3 | 0.460±0.009 | 0.323±0.009 | 0.22±0.02 |
| | DMPC | n=6 | 0.46±0.04 | 0.30±0.03 | 0.24±0.03 |
| $ApoE4_{182, 241}$ | lipid-free | n=4 | 0.22±0.01 | 0.57±0.03 | 0.21±0.02 |
| | DMPC | n=4 | 0.29±0.01 | 0.50±0.02 | 0.21±0.02 |
| $ApoE4_{223, 291}$ | lipid-free | n=4 | 0.13±0.02 | 0.77±0.04 | 0.11±0.02 |
| | DMPC | n=3 | 0.227±0.005 | 0.62±0.02 | 0.16±0.02 |
| $ApoE4_{86, 241}$ | lipid-free | n=3 | 0.131±0.006 | 0.79±0.02 | 0.08±0.01 |
| | DMPC | n=3 | 0.20±0.02 | 0.71±0.03 | 0.09±0.01 |

**Supplementary Table 15. Analysis of Fluorescence Correlation Spectroscopy under lipid-bound conditions.**

| Lipid-bound (DMPC) ApoE4 constructs | | | | | |
|---|---|---|---|---|---|
| **Construct** | **$R_h^{DD}$(nm)** | **$R_h^{AA}$(nm)** | **$N^{DD}$** | **$N^{AA}$** | |
| $ApoE4_{5,\ 86}$ | 6.3±0.7 | 9±1 | 0.4±0.1 | 0.13±0.02 | n=3 |
| $ApoE4_{86,\ 165}$ | 9±2 | 10±2 | 0.25±0.04 | 0.097±0.003 | n=6 |
| $ApoE4_{182,\ 241}$ | 9±1 | 9.2±0.5 | 0.3±0.1 | 0.100±0.006 | n=4 |
| $ApoE4_{223,\ 291}$ | 11.5±0.4 | 14±2 | 0.27±0.04 | 0.109±0.009 | n=4 |
| $ApoE4_{86,\ 241}$ | 12±1 | 12±3 | 0.212±0.009 | 0.17±0.04 | n=3 |

**Supplementary Table 16. Residue pairs used for computing distances used in clustering protein conformations.**

| Residue pairs | | |
|---|---|---|
| 5,86 | 48,129 | 32,67 |
| 86,165 | 67,149 | 108,149 |
| 182,241 | 129,234 | 32,149 |
| 223,291 | 32,108 | 67,108 |
| 86,241 | 191,215 | 11,280 |
